# Bayesian estimation for stochastic gene expression using multifidelity models

**DOI:** 10.1101/468090

**Authors:** Huy D. Vo, Zachary Fox, Ania Baetica, Brian Munsky

## Abstract

The finite state projection (FSP) approach to solving the chemical master equation has enabled successful inference of discrete stochastic models to predict single-cell gene regulation dynamics. Unfortunately, the FSP approach is highly computationally intensive for all but the simplest models, an issue that is highly problematic when parameter inference and uncertainty quantification takes enormous numbers of parameter evaluations. To address this issue, we propose two new computational methods for the Bayesian inference of stochastic gene expression parameters given single-cell experiments. We formulate and verify an Adaptive Delayed Acceptance Metropolis-Hastings (ADAMH) algorithm to utilize with reduced Krylov-basis projections of the FSP. We then introduce an extension of the ADAMH into a Hybrid scheme that consists of an initial phase to construct a reduced model and a faster second phase to sample from the approximate posterior distribution determined by the constructed model. We test and compare both algorithms to an adaptive Metropolis algorithm with full FSP-based likelihood evaluations on three example models and simulated data to show that the new ADAMH variants achieve substantial speedup in comparison to the full FSP approach. By reducing the computational costs of parameter estimation, we expect the ADAMH approach to enable efficient data-driven estimation for more complex gene regulation models.

## INTRODUCTION

An important goal of quantitative biology is to elucidate and predict the mechanisms of gene expression. Evidence increasingly suggests that gene expression processes are inherently stochastic with substantial cell-to-cell variability. ^1–3^ In an isogenic population with the same environmental factors, much of these fluctuations can be attributed to intrinsic chemical noise. There are different experimental methods to obtain information about the stochastic behavior of single cells,^4^ each producing a unique type of data that necessitates a different statistical and computational framework to interpret the observed gene expression dynamics.^5^ Our present work focuses specifically on building Bayesian computational tools to analyze and model data from smFISH and related techniques.^6–10^ A sucessful framework for building predictive models for gene expression dynamics from such data is to fit the solution of the chemical master equation (CME)^11^ to the empirical histogram obtained from population snapshots at several experimental conditions or time-points. ^8,10,12,13^

The finite state projection (FSP),^14^ which approximates the dynamics of the CME with a finite system of linear ODEs, provides a framework to analyze full distributions of stochastic gene expression models with computable error bounds. It has been observed that full distribution-based analyses using the FSP perform well, even when applied to realistically small experimental datasets on which summary statistics-based fits may fail. ^15^ On the other hand, the FSP requires solving a large system of ODEs that grows quickly with the complexity of the gene expression network under consideration. Our present study borrows from model reduction strategies in other complex systems fields to alleviate this issue by reducing the computational cost of FSP-based parameter estimation.

There has been intensive research on efficient computational algorithms to quantify the uncertainty in complex models.^16^ A particularly promising approach is to utilize multifidelity algorithms to systematically approximate the original system response. In these approximations, surrogate models or meta-models allow for various degrees of model fidelity (e.g., error compared to the exact model) in exchange for reductions in computational cost. Surrogate models generally fall into two categories: response surface and low-fidelity models. ^17,18^ We will focus on the second category that consists of reduced-order systems, which approximate the original high-dimensional dynamical system using either simplified physics or projections onto reduced order subspaces. ^16,19,20^ Reduced-order modeling has already begun to appear in the context of stochastic gene expression. When all model parameters are known, the CME can be reduced by system-theoretic methods, ^21,22^ sparse-grid/aggregation strategies, ^23,24^ tensor train representations^25–27^ and hierarchical tensor formats.^28^ Model reduction techniques have also been applied to parameter optimization by Waldherr and Hassdonk ^29^ who projected the CME onto a linear subspace spanned by a reduced basis, and Liao et al. ^30^ who approximated the CME with a Fokker-Planck equation that was projected onto the manifold of low-rank tensors. ^31^ While these previous works clearly show the promise of reduced-order modeling, there remains a vast reservoir of ideas from the broader computational science and engineering community that remain to be adapted to the quantitative analysis of stochastic gene expression.

In this paper, we introduce two efficient algorithms, which are based on the templates of the adaptive Metropolis algorithm^32^ and the delayed acceptance Metropolis-Hastings (DAMH^33,34^) algorithm, to sample the posterior distribution of gene expression parameters given single-cell data. The adaptive Metropolis approach automatically tunes parameter proposal distributions to more efficiently search spaces of unnormalized and correlated parameters. The DAMH provides a two-stage sampling approach that uses a cheap approximation to the posterior distribution at the first stage to quickly filter out proposals with low posterior probabilities. Improvements to the DAMH allow algorithmic parameters to be updated adaptively and automatically by the DAMH chain. ^35,36^ The DAMH has been applied to the inference of stochastic chemical kinetics parameters from time-course data in combination with approximate particle filtering schemes based on the Chemical Langevin equation (CLE) and the Linear Noise approximation (LNA). ^37^ Our algorithm is a modified version of DAMH that is specifically adapted to improve Bayesian inference from specific-time snapshots of relatively small populations of single cells, such as that which arises from smFISH and other optical microscopy experiments. We employ parametric reduced order models using Krylov-based projections,^38,39^ which give an intuitive means to compute expensive FSP-based likelihood evaluations. ^40,41^ To improve the accuracy and the DAMH acceptance rate, we allow the reduced model to be refined during parameter space exploration. The resulting method, which we call the ADAMH-FSP-Krylov algorithm, is tested on three common gene expression models. We also provide a theoretical guarantee and numerical demonstrations that the proposed algorithms converge to equivalent target posterior distributions.

The organization of the paper is as follows. We review the background on the FSP analysis of single-cell data, and basic Markov chain Monte Carlo (MCMC) schemes in the *Background* section. In the *Materials and Methods* section, we introduce our method to generate reduced FSP models, as well as our approach to monitor and refine their accuracy. These reduced models provide an approximation to the true likelihood function, which is then employed to devise an Adaptive Delayed Acceptance Metropolis-Hastings with FSP-Krylov reduced models (ADAMH-FSP-Krylov) and a Hybrid algorithm. We make simple adjustments to the existing ADAMH variants in the literature to prove convergence, and we give the mathematical details in the supplementary materials. We provide empirical validation of our methods on three gene expression models using synthetic data sets, and we compare the efficiency and accuracy of the approaches in the *Results* section. Interestingly, we find empirically that the reduced models learned through the ADAMH run could fully substitute the original FSP model in a Metropolis-Hastings run without incurring a large difference in the sampling results. Finally, we conclude with a discussion of future work and the potential of computational science and engineering tools to analyze stochastic gene expression.

## BACKGROUND

### Stochastic modeling of gene expression and the chemical master equation

Consider a well-mixed biochemical system with *N* ≥ 1 different chemical species that are interacting via *M* ≥ 1 chemical reactions. Assuming constant temperature and volume, the time-evolution of this system can be modeled by a continuous-time Markov process. ^11^ The state space of the Markov process consists of integral vectors ***x*** ≡ (*x*_1_,…, *x*_*N*_) ^*T*^, where *x*_*i*_ is the population of the *i*th species. Each reaction channel, such as the transcription of an RNA species, is characterized by a *stoichiometric* vector ***v***_*j*_ (*j* = 1,…, *M*) that represents the change when the reaction occurs; if the system is in state ***x*** and reaction *j* occurs, then the system transitions to state ***x*** + ***v***_*j*_. Given ***x***(*t*) = ***x***, the propensity *α*_*j*_(***x***; ***θ***)*dt* determines the probability that reaction *j* occurs in the next infinitesimal time interval [*t, t* + *dt*), where ***θ*** is the vector of model parameters.

Since the state space is discrete, we can index the states as ***x***_1_,…, ***x***_*n*_,…. The time-evolution of the probability distribution of the Markov process is the solution of the linear system of differential equations known as the chemical master equation (CME):

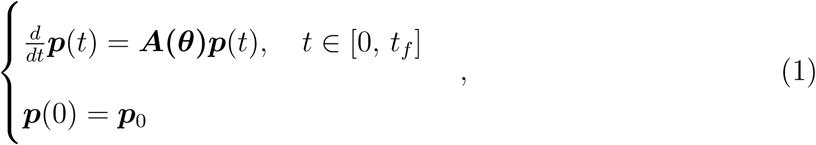

where the probability mass vector ***p*** = (*p*_1_, *p*_2_,…)^*T*^ is such that each component, *p*_*i*_ = *P* (*t*, ***x***_*i*_) = Prob{***x***(*t*) = ***x***_*i*_}, describes the probability of being at state ***x***_*i*_ at time *t*, for *i* = 1,…, *n*. The vector ***p***_0_ = ***p***(0) is an initial probability distribution, and ***A***(***θ***) is the infinitesimal generator of the Markov process. Here, we have made explicit the dependence of ***A*** on the model parameter vector ***θ***, which is often inferred from experimental data.

### Finite State Projection

The state space of the CME could be infinite or extremely large. To alleviate this problem, the finite state projection (FSP^14^) was introduced to truncate the state space to a finite size. In the simplest FSP formulation, the state space is restricted to a hyper-rectangle

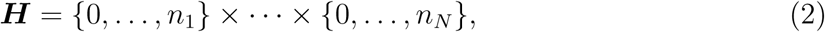

where the *n*_*k*_ are the maximum copy numbers of the chemical species.

The infinite-dimensional matrix ***A*** and vector ***p*** in eq. (1) are replaced by the corresponding submatrix and subvector. When the bounds *n*_*k*_ are chosen sufficiently large and the propensities satisfy some regularity conditions, the gap between the FSP and the original CME is negligible and computable.^14,42^ Throughout this paper, we assume that the bounds *n*_*k*_ have been chosen appropriately and that the FSP serves as a high-fidelity model of the gene expression dynamics of interest. Our goal is to construct lower-fidelity models of the FSP using model order reduction and incorporate these reduced models in the uncertainty analysis for gene expression parameters.

### Bayesian inference from single-cell data

Data from smFISH experiments ^6–8,10^ consist of several snapshots of many independent cells taken at discrete times *t*_1_,…, *t*_*T*_. The snapshot at time *t*_*i*_ records gene expression in *n*_*i*_ cells, each of which can be collected in the data vector ***c***_*j,i*_, *j* = 1,…, *n*_*i*_ of molecular populations in cell *j* at time *t*_*i*_. Let ***p***(*t*, ***x***|***θ***) denote the entry of the FSP solution corresponding to state ***x*** at time *t*, with model parameters ***θ***. The FSP-based approximation to the log-likelihood of the data set 𝒟 given parameter vector ***θ*** is given by

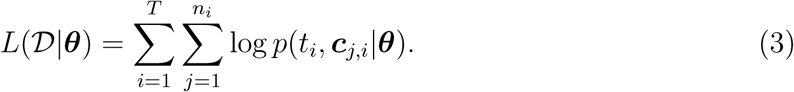

It is clear that when the FSP solution converges to the true solution of the CME, the FSP-based log-likelihood converges to the true data log-likelihood. The posterior distribution of model parameters ***θ*** given the data set 𝒟 then takes the form

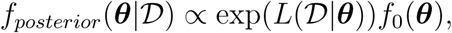

where *f*_0_ is the prior density that quantifies prior knowledge and beliefs about the parameters. When *f*_0_ is a constant, the parameters that maximize the posterior density are equivalent to the maximum likelihood estimator. However, we also want to quantify our uncertainty regarding the accuracy of the parameter fit, and the MCMC framework provides a way to address this by sampling from the posterior distribution.

For convenience, we limit our current discussion to models and inference problems that have the following characteristics:

1. The matrix ***A***(***θ***) can be decomposed into

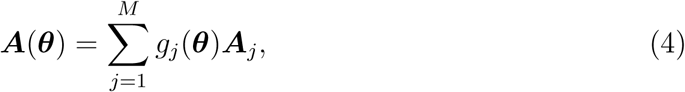

where *g*_*j*_ are continuous functions and ***A***_*j*_ are independent of the parameters.

2. The support of the prior is contained in a bounded domain of the form

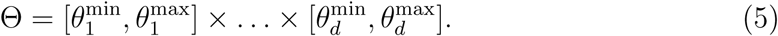

The first assumption means that the CME matrix depends “linearly” on the parameters, ensuring the efficient assembly of the parameter-dependent matrix. In particular, the factors ***A***_*j*_ can be computed and stored in the offline phase before parameter exploration and only a few (sparse) matrix additions are required to compute ***A***(***θ***) in the online phase. When there are nonlinear dependence on parameters, more sophisticated methods such as the Discrete Empirical Interpolation method ^43^ could be applied, but we leave this development for future work in order to focus more on the parameter sampling aspect. Nevertheless, condition (4) covers an important class of models, including all models defined by mass-action kinetics. The second assumption means that the support of the posterior distribution is a bounded and well-behaved domain (in mathematical terms, a compact set). This allows us to derive convergence theorems more straightforwardly. In practice, condition (5) is not a severe restriction since it can be interpreted as the prior belief that physical parameters cannot assume infinite values.

### The Metropolis-Hastings and the adaptive Metropolis algorithms

The Metropolis-Hastings (MH) Algorithm ^44,45^ is one of the most popular methods to sample from a multivariate probability distribution (Algorithm 1). The basic idea of the MH is to generate a Markov chain whose limiting distribution is the target distribution. To do so, the algorithm includes a probabilistic acceptance/rejection step. More precisely, let *f* denote the target probability density. Assume the chain is at state ***θ***_*i*_ at step *i*. Let ***θ*′** be a proposal from the pre-specified proposal density *q*(.|***θ***_*i*_). The DAMH computes a first-step acceptance probability of the form

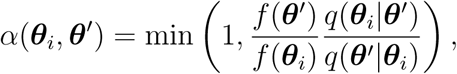

to decide whether to accept ***θ*′** as the next state of the chain. If ***θ* ′** fails to be promoted, the algorithm moves on to the next iteration with ***θ***_*i*+1_ := ***θ***_*i*_.

There could be many choices for the proposal density *q* (for example, see the survey of Roberts and Rosenthal^46^). We will consider only the symmetric case where *q* is a Gaussian, that is,

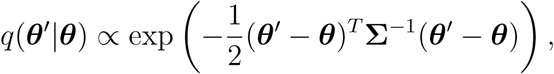

where **Σ** is a positive definite matrix that determines the covariance of the proposal distribution. With this choice of a symmetric proposal distribution, the MH reduces to the original Metropolis Algorithm. ^44^ For gene expression models, the MH has been combined with the FSP for parameter inference and model selection in several studies. ^10,15^

The appropriate choice of **Σ** is crucial for the performance of the Metropolis algorithm. Haario et al. ^32^ proposes an Adaptive Metropolis (AM) algorithm in which the proposal **Σ** is updated at every step using the values visited by the chain. This is the version that we will implement for sampling the posterior distribution with the full FSP model. In particular, let ***θ***_1_,…, ***θ***_*i*_ be the samples accepted so far, the AM updates the proposal covariance using the formula

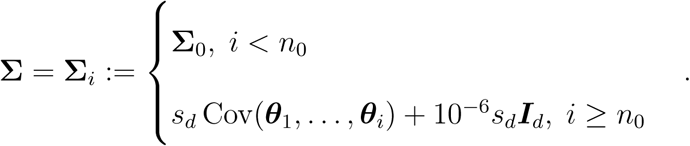

Here, the function Cov returns the sample covariances. The constant s_d_ is assigned the value (2.4)^2^*/d* following Haario et al.^32^ The matrix **Σ**_0_ is an initial choice for the Gaussian proposal density, and *n*_0_ is the number of initial steps without proposal adaptations. Using the adaptive Metropolis allows for more efficient search over un-normalized and correlated parameters spaces and eliminates the need for the user to manually tune the algorithmic parameters. In the numerical results that we will show, the adaptive Metropolis results in reasonable acceptance rates (19% - 23.4%). The adaptive MH has been used in previous works on gene expression models in combination with fluorescent time-course data and flow cytometry data.^47,48^

## MATERIALS AND METHODS

### Delayed acceptance Metropolis-Hastings algorithm

Previous applications of the MH to gene expression have required 10^4^ to 10^6^ or more iterations per combination of model and data set,^15^ and computational cost is a significant issue when sampling from a high-dimensional distribution whose density is expensive to evaluate. A practical rule of thumb for balancing between exploration and exploitation for a MH algorithm with the Gaussian proposal is to have an acceptance rate close to 0.234, which was derived by Roberts et al.^49^ as the asymptotically optimal acceptance rate for random walk MH algorithms. Assuming the proposal density of Algorithm 1 is tuned to have an acceptance rate of approximately 23.4%, one could achieve significant improvement to computation time by quickly rejecting poor proposals without evaluating the expensive posterior density.

#### Algorithm 1

Metropolis-Hastings

**Input:**

Target density *f* (.);

Initial parameter ***θ***_0_;

Proposal density *q*(*·*|*·*);

1: **for** *i* = 0, 1,…, **do**

2: Draw ***θ***^*1*^ from the proposal density *q*(***θ***_*i*_)

3: Compute the acceptance probability

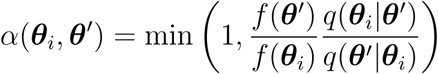

4: With probability *α*(***θ***_*i*_, ***θ*′**), set ***θ***_*i*+1_←***θ*′** (accept); otherwise ***θ***_*i*+1_ ←***θ***_*i*_ (reject).

5: **end for**

**Output:** samples ***θ***_1_, ***θ***_2_,…

The delayed acceptance Metropolis-Hasting (DAMH)^33^ seeks to alleviate the computational burden of rejections in the original MH by employing a rejection step based on a cheap approximation to the target density (cf. Algorithm 2). Specifically, let *f* (.) be the density of the target distribution of the parameter ***θ***. Let 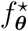(.) be a cheap state-dependent approximation to *f*. At iteration *i*, let ***θ*′** be a proposal from the current parameter ***θ*** using a pre-specified proposal density *q*(.|.). The DAMH promotes ***θ′*** as a potential candidate for acceptance with probability

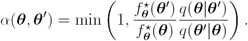

If ***θ*′** fails to be promoted, the algorithm moves on to the next iteration with ***θ***_*i*+1_ := ***θ***_*i*_. If the ***θ′*** passes the first inexpensive check, than a second acceptance probability is computed using the formula

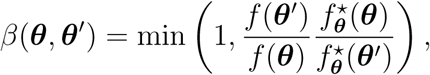

and the DAMH algorithm accepts ***θ′*** for the next step with probability *β*. In this manner, computational savings can be expected if unlikely proposals are quickly rejected in the first step, leaving only the most promising candidates for careful evaluation in the second step. Christen and Fox show that the ADAMH converges to the target distribution under conditions that are easily met in practice.^33^ However, the quality of the approximation 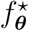 affects the overall efficiency. Poor approximations lead to many false promotions of parameters that are rejected at the expensive second step. On the other hand, the first step may falsely reject parameters that could have been accepted using the accurate log-likelihood evaluation. This leads to subsequent developments that seek appropriate approximations and ways to adapt these approximations to improve the performance of DAMH in specific applications. ^35,50^ Specifically, the adaptive DAMH variant in Cui et al., 2014^50^ formulates 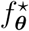 via reduced basis models that can be updated on the fly using samples accepted by the chain. The adaptive version in Cui et al., 2011, ^35^ allows adaptations for the proposal density and the error model, with convergence guarantees. ^36^ We will borrow these elements in our sampling scheme that we introduce below. However, the stochastic gene expression models that we investigate here differ from the models studied in those previous contexts, since our likelihood function incorporates intrinsic discrete state variability instead of external Gaussian noise.

## Reduced-order models for the FSP dynamics

### Projection-based parametric model reduction

#### Algorithm 2

Delayed Acceptance Metropolis-Hastings

**Input:**

Target density *f* (.);

State-dependent density *approximation* 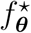(.);

Initial parameter ***θ***_0_;

Proposal density *q*(*·|·*);

1: **for** *i* = 0, 1, *…*, **do**

2: Draw ***θ′*** from the proposal density *q*(·|***θ***_*i*_)

3: Compute the first-stage acceptance probability

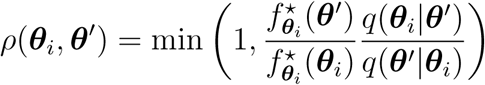

4: With probability *ρ*(***θ***_*i*_, ***θ′***), promote the value of ***θ′*** to the next stage. Otherwise, set ***θ*** _*i+1*_, ← ***θ***_*i*_.

5: If ***θ′*** was promoted, compute the second-stage acceptance probability

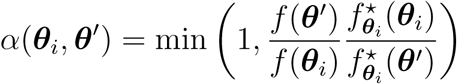

6: With probability *α*(***θ***_*i*_, ***θ′***), set ***θ***_*i*+1_ *←* ***θ′*** (accept); otherwise ***θ***_*i*+1_*←****θ***_*i*_ (reject).

7: **end for**

**Output:** samples ***θ***_1_, ***θ***_2_, *…*

In this subsection, we review the principle of projection-based model reduction, which consists of projecting a high-dimensional dynamics onto a low-dimensional subspace. ^20^ In particular, consider the parameter-dependent FSP dynamics

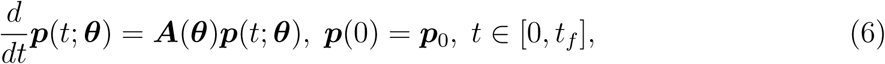

where the parameter-dependent transition rate matrix ***A***(***θ***) satisfies assumption (4) and ***p***(*t*; ***θ***) is the high-fidelity approximation to the probability distribution of the CME. Consider a user-specified partitioning of the time interval [0, *t*_*f*_] into *n*_*B*_ subintervals [*t*_*i-*1_, *t*_*i*_] with *i* = 1, *…, n*_*B*_ and *t*_0_ = 0. We will sequentially project the high-fidelity dynamical system onto a sequence of low-dimensional subspaces associated with these subintervals. In particular, consider an ordered set of reduced bases 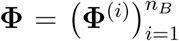, where 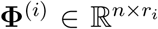, *r*_*i*_ *≤ n* is the orthogonal basis matrix of the subspace associated with the *i*-th subinterval. We seek an approximation of the form

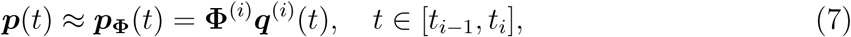

which leads to the approximate dynamical system

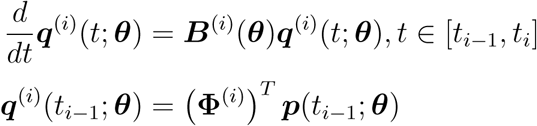

where ***B***^(*i*)^(***θ***) = (**Φ**^(*i*)^)^*T*^ ***A***(***θ***)**Φ**^(*i*)^. Under assumption (4), the reduced system matrices ***B***^(*i*)^(***θ***) in eq. (9) can be decomposed as

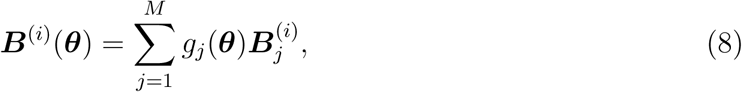

where 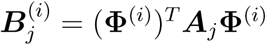. This decomposition allows us to assemble the reduced systems quickly with 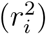 complexity. Using the approximation ***p***(*t*_*i-*1_; ***θ***) ≈ **Φ**^(*i-*1)^***q***^(*i-*1)^(*t*_*i-*1_) again and substitute this to the preceding reduced system, we get

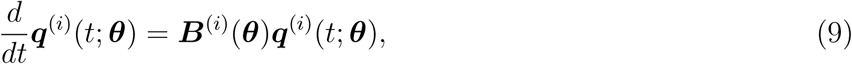

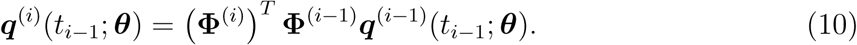

This results in a sequence of reduced-order models that we will use for our model reduction method. Equation (10) implies that the solution at a previous time interval will be projected onto the subspace of the next interval. While this introduces some extra errors, subdividing the long time interval helps to reduce the subspace dimensions for systems with complicated dynamics.

The choice of the basis set **Φ** clearly determines the approximation quality of the reduced systems. Our basis construction procedure proceeds at two levels. First, we construct local bases that yield approximate models of the FSP at individual parameters. From there, we proceed to construct a global basis that approximates the FSP model over the whole parameter domain. The details of this procedure are explained in the next two subsections.

### Krylov subspace approximation for single-parameter model reduction

Consider a fixed parameter combination ***θ***. Let the time points 0 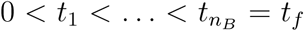 be given. Using a high-fidelity solver, we can compute the full solution at those time points, and we let ***p***_*i*_ denote the full solution at time *t*_*i*_. Our aim is to construct a sequence of orthogonal matrices ***V*** ^(*i*)^ *≡* ***V*** ^(*i*)^(***θ***) with *i* = 1 *… n*_*B*_ such that the full model dynamics at the parameter ***θ*** on the time interval [*t*_*i−*1_, *t*_*i*_] is well-approximated by a projected reduced model on the span of ***V*** ^(*i*)^.

A simple and effective way to construct the reduced bases is to choose ***V*** ^(*i*)^ as the orthogonal basis of the Krylov subspace

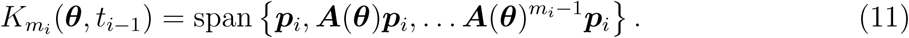

Here, *𝒱* = span *U* means that every element of *𝒱* is a linear combination of a finite number of elements in *U*. In order to determine the subspace dimension *m*_*i*_, we use the error series derived by Saad^38^ which we reproduce here using our notation as

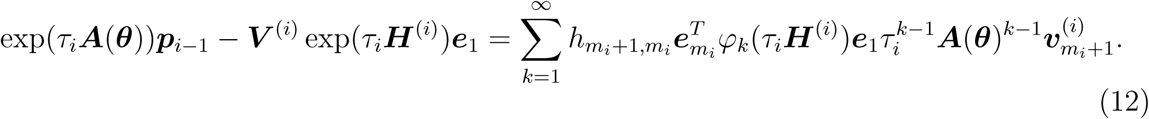

Here, 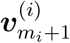 and 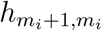 are the outputs at step *m*_*i*_ of the Arnoldi procedure (Algorithm10.5.1 in Golub and Van Loan^51^) to build the orthogonal matrix ***V*** ^(*i*)^, where 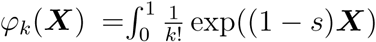 ds for any square matrix ***X***. The matrix ***H***^(*i*)^ = (***V*** ^(*i*)^)^*T*^ ***A*(*θ*)*V*** ^(*i*)^ is the state matrix of the reduced-order system obtained via projecting ***A*** onto the Krylov subspace *K*_*m*_. The terms 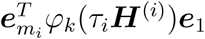 can be computed efficiently using Expokit (Theorem 1, Sidje^39^). We use the Euclidean norm of the first term of the series (12) as an indicator for the model reduction error. Given an error tolerance *ε*_Krylov_, we iteratively construct the Krylov basis ***V*** ^(*i*)^ with increasing dimension until the error per unit time step of the reduced model falls below the tolerance, that is,

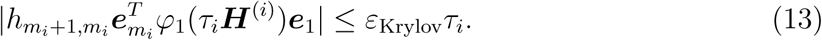

We clarify that the Krylov-based model reduction method has been around for decades and has been applied to the solution of linear systems many non-biological fields. Krylov subspaces provide a way to integrate the full FSP model in a non-parameteric setting, as studied in many previous works.^40,41,52,53^ The novelty of our work is in using the Krylov bases constructed at different parameters in order to produce a global basis that approximates the parameter-dependent FSP over the entire parameter domain. This is explained below.

### Global basis construction

We build the reduced basis for the parameter-dependent dynamics by concatenation (see, e.g., Benner et al. ^20^). Specifically, for any ***θ*** let 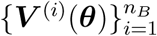 be the sequence of orthogonal basis matrices constructed with Krylov projection as described above. We can sample different bases from a finite set of ‘training’ parameters 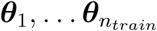. Then, through the iterative updates

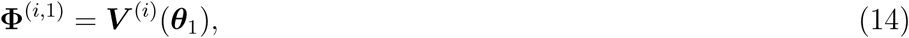

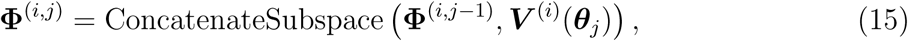

we obtain the bases 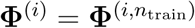 that provide global approximations for the full dynamical system across the parameter domain. The operator ConcatenateSubspace(***X***, ***Y***), assuming ***X*** is already orthogonal, builds an orthogonal basis matrix for the union of the subspaces spanned by the columns of both ***X*** and ***Y*** by first orthogonalizing each column of ***Y*** against all columns of ***X*** and concatenating these orthogonalized columns to ***X***.

The selection of the training instances 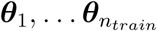 influences the quality of the global bases. The classical Greedy scheme^54^ consists of an *offline* and an *online* phase. All training parameters are generated from the prior distribution in the offline phase and a global basis is constructed from these training parameters. In the online phase, only the reduced model is used for optimization and other purposes. However, since the posterior is necessarily shaped differently than the prior, building a global basis based on the prior distribution is inefficient for the purpose of MCMC analysis. In particular, the offline phase may sample from from regions with low posterior density, thus adding excessive information that is not needed by the MCMC. We instead follow the approach of Cui et al.,^50^ which builds a small initial basis using the MCMC starting poiint and update this basis as the chain explores more parameters. This is accomplished via the Delayed Acceptance framework. We give more details below.

## Adaptive Delayed Acceptance Metropolis with reduced-order models of the CME

### The approximate log-likelihood formula

Following the discussion above, suppose that an initial reduced basis **Φ** is given, and we have a reduced-cost approximations ***p*** ≈ ***p***_**Φ**_ to the full FSP dynamics. We can then approximate the full log-likelihood of single-cell data in equation (3) by the reduced-model-based log-likelihood

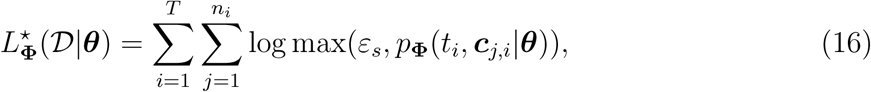

where *ε*_*s*_ is a small constant, chosen to safeguard against undefined values. We need to include *ε*_*s*_ in our approximation since the entries of the reduced-order approximation are not guaranteed to be positive (not even in exact arithmetic). We aim to make the approximation to be accurate for parameters ***θ*** with high posterior density, and crude on those with low density, which should be visited rarely by the Monte Carlo chain.

One can readily plug in the approximation (16) to the DAMH algorithm. Since 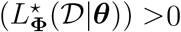 for all 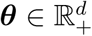, the chain will eventually converge to the target posterior distribution (Theorem 1 in Christen and Fox, ^33^ and Theorem 3.2 in Efendiev et al. ^34^). On the other hand, a major problem with the DAMH is that the computational efficiency depends on the quality of the reduced basis approximation. Crude models result in high rejection rates at the second stage, thus increasing sample correlation and computation time. Therefore, it is advantageous to fine-tune the parameters of the algorithm and update the reduced models adaptively to ensure a reasonable acceptance rate. This motivates the adaptive version of the DAMH, which we discuss next.

### Delayed acceptance posterior sampling with infinite model adaptations

We propose an adaptive version of the DAMH for sampling from the posterior density of the CME parameters given single-cell data (Algorithm 3). We have borrowed elements from the adaptive DAMH algorithms in Cui et al.^35,50^ The first step proposal uses an adaptive Gaussian similar to the adaptive Metropolis of Haario et al.,^32^ where the covariance matrix is updated at every step from the samples accepted so far.

The reduced bases are updated as the chain explores the parameter domain. Instead of using a finite adaptation criterion to stop model adaptation as in Cui et al., ^50^ we introduce an adaptation probability with which the reduced basis updates are considered. This means that an infinite amount of model adaptations could occur with diminishing probability as the chain progresses. This idea is taken from the “doubly-modified example” in Roberts and Rosenthal. ^55^ The advantage of the probabilistic adaptation criteria is that it allows us to prove ergodicity for the adaptive algorithm. The mathematical proofs are presented in the Supportiing Information.

The adaptation probability *a*(*i*) is chosen to converge to 0 as the chain iteration index *i* increases. In particular, we use

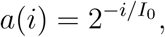

where *I*_0_ is a user-specified constant. This formula states that the probability for an adaptation to occur decreases by half after every *I*_0_ chain iterations. In addition, we further restrict the adaptation to occur only when the error indicator is above a threshold at the proposed parameters. As a consequence of our model updating criteria, the reduced-order bases will be selected at points that are close to the support of the target posterior distribution.

#### Algorithm 3

ADAMH-FSP-Krylov

**Input:**

Prior density *f*_0_;

Parameter-dependent CME matrix ***A***(.);

Chain starting point ***θ***_0_, initial proposal covariance ***C***_0_.

Basis update tolerance *ε*_*basis*_, Krylov tolerance *ε*_*Krylov*_, reduced model time partition 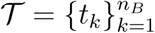;

Adaptation probability 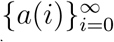;

Maximum basis dimension *m*_*max*_.

1: **Φ**_0_ = GenerateKrylovBases(***A***(***θ***_0_), ***p***_0_, *𝒯, ε*_*Krylov*_);

2: **for** *i* = 0, 1,*…* **do**

3: Compute the proposal *θ′* ∼ *N* (***θ***_*i*_, ***C***_*i*_);

4: With probability *α*, promote *θ′*, where

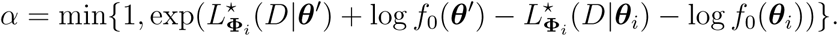

5: Otherwise, ***θ***_*i*+1_ := ***θ***_*i*_ and move on to the next iteration.

6: **if** *θ′* was promoted **then**

7: With probability *β*, accept *θ′* as the next sample ***θ***_*i*+1_. Here,

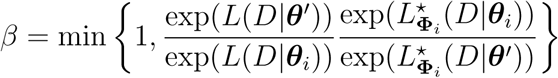

8: Otherwise, set ***θ***_*i*+1_ = ***θ***.

9: Compute ErrorEst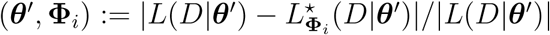

10: **if** ErrorEst(*θ′*, **Φ**_*i*_) *> ε*_*basis*_ and *θ′* was accepted **then**

11: With probability *a*(*i*), **Φ**_*i*+1_ = UpdateBases(**Φ**_*i*_, ***θ***_*i*_), otherwise **Φ**_*i*+1_ := **Φ**_*i*_.

12: **else**

13: **Φ**_*i*+1_ := **Φ**_*i*_.

14: **end if**

15: **end if**

16: 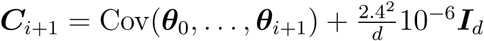 ▹Update the proposal density

17: **end for**

**Output:**Samples ***θ***_0_, ***θ***_1_,*…*

## Using ADAMH in approximate Bayesian inference

A limitation of the delayed acceptance schemes we presented above is that every accepted parameter incurs an expensive FSP evaluation. Trivially, the number of full model solutions could be reduced by using a proposal with low acceptance rate. However, this leads to statistical inefficiency and eventually we still have to run very long chains to obtain enough independent samples. In many realistic situations, we may only be able to solve the full models for a limited number of times. Drawing from an approximate posterior distribution becomes the only viable option. The ADAMH provides a way to construct such an approximate posterior. In particular, we propose a ‘Hybrid’ chain that spends the first few iterations using the same transition kernel as the ADAMH. These first steps allow us to gradually build up a reduced basis to approximate the true FSP model across parameters with high posterior probability densities. Once a sufficiently good model has been constructed (or we have exceeded the maximum budget allowed for the full FSP solves), the chain transits to a second phase in which only the cheap approximations provided by the reduced model are used for the Metropolis-Hastings criteria. We test a prototypical implementation of this idea in the next section, where we found that using the two-phase scheme can bring dramatic speedup (up to an order of magnitude for a sizeable example) while giving almost identical results to the adaptive Metropolis-Hastings and ADAMH schemes. A similar strategy was previously introduced by Cui et al.^50^ for inverse problems related to elliptic PDEs.

## RESULTS

We conduct numerical tests on several stochastic gene expression models to study performance of our proposed Algorithms. For the first two examples, the test platform is a desktop computer running Linux Mint and MATLAB 2018b, with 32 GB RAM and Intel Core i7 3.4 GHz quad-core processor. The last example is tested on a single node of the Keck Computing Cluster at Colorado State University, which consists of 16 cores Xeon E5-2620 v4, with 64 GB RAM with MATLAB 2017b installed. Both MATLAB versions are allowed to use the maximum number of threads available on each machine, which are respectively four (Desktop) and 16 (cluster node).

We compare three sampling algorithms:

1. Adaptive Metropolis-Hastings with full FSP-based likelihood evaluations (AMH-FSP): This version is an adaptation of the Adaptive Metropolis of Haario et al., ^32^ which updates the covariance of the Gaussian proposal density at every step. The algorithm always uses the FSP-based likelihood (3) to compute the acceptance probability, and it is solved using the Krylov-based Expokit. ^39^ This is the reference algorithm by which we assess the accuracy and performance of the other sampling schemes. This is the best scheme that we know of that can automatically balance between exploration and exploitation for the MCMC samples. The AMH has been used in several previous works to investigate gene expression models.^47,48^
2. Adaptive Delayed Acceptance Metropolis-Hastings with reduced FSP model constructed from Krylov subspace projections (ADAMH-FSP-Krylov): This is Algorithm 3 mentioned above. Similar to AMH-FSP, this algorithm uses a Gaussian proposal with an adaptive covariance matrix. However, it has a first-stage rejection step that employs the reduced model constructed adaptively using Krylov-based projection.
3. Hybrid method that consists of two phases. A reduced model is constructed during the first phase as in the ADAMH-FSP-Krylov. The second phase then uses the Adaptive Metropolis-Hastings updates with only reduced model-based likelihood evaluations. This is similar to AMH-FSP, but we instead use the approximate log-likelihood formula (16). For a user-specified chain length *𝓁*, we let the first phase persist for the first ten percent of the iterations, and switch to the second phase for the rest of the iterations. The ten percent threshold is tentative, and it comes from our observation that the ADAMH-FSP-Krylov performs most of its model updates during the first ten percent of the iterations. This scheme does not have strong asymptotic convergence guarantees as the ADAMH-FSP-Krylov, but we will see from the numerical tests that it can still yield similar parameter estimation results to the other schemes in dramatically less computation time.

We rely on two metrics for performance evaluation: total CPU time to finish each chain, and the multivariate effective sample size as formulated in Vats et al. ^56^ Given samples ***θ***_1_, *…*, ***θ***_*n*_, the multivariate effective sample size is estimated by

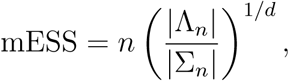

where Λ_*n*_ is an estimation of the posterior covariance using the sample covariance, and Σ_*n*_ the multivariate batch means estimator. An algorithm, whose posterior distribution matches the full FSP implementation, but with a lower ratio of CPU time per (multivariate) effective sample will be deemed more efficient. We use the MATLAB implementation by Luigi Acerbi ^1^ for evaluating the effective sample size from the MCMC outputs. We also conduct Geweke diagnostics^57^ and calculate the integrated autocorrleation time (IACT) of the chains using the MATLAB package MCMCSTAT ^2^.

In all examples considered, the prior and the parameters are first transformed into log10 scale before applying the MCMC algorithms. This transformation allowed a better exploration of the parameter space, especially for cases where the parameters are poorly constrained by the data.

To achieve reproducible results for each example, we reset the random number generator to Mersenne Twister with seed 0 in Matlab before simulating the single-cell observations with Gillespie’s Algorithm ^58^ and running the three algorithms for a specified number of iterations.

### Two-state gene expression

We first consider the common model of bursting gene expression ^7,59–62^ with a gene that can switch between ON and OFF states and an RNA species that is transcribed when the gene is switched on (Table 1).

**Table 1:** Two-state gene expression reactions and propensities.

| reaction | propensity |
| --- | --- |
| 1. $G_{OFF} \xrightarrow{k_{on}} G_{ON}$ | $\alpha_1 = k_{on}[G_{OFF}]$ |
| 2. $G_{ON} \xrightarrow{k_{off}} G_{OFF}$ | $\alpha_2 = k_{off}[G_{ON}]$ |
| 3. $G_{ON} \xrightarrow{k_r} G_{ON} + RNA$ | $\alpha_3 = k_r[G_{ON}]$ |
| 4. $RNA \xrightarrow{\gamma} \emptyset$ | $\alpha_4 = \gamma[RNA]$ |
| Parameters' units are $\text{hr}^{-1}$. $[X]$ is the number of copies of the species X. | |

We simulate data at ten equally spaced time points from 0.1 to 1 hour, with 200 independent observations per time point. The gene states are assumed to be unobserved. We generate the reduced bases on subintervals generated by the time points in the set

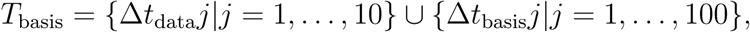

where Δ*t*_data_ = 0.1hr and Δ*t*_basis_ = 0.01hr. Thus, *T*_basis_ includes the observation times. We choose the basis update threshold as *δ* = 10^−4^. The prior distribution in our test is the log-uniform distribution on a rectangle, whose bounds are given in Table 2. The full FSP state space is chosen as

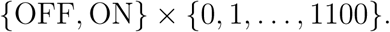

We choose a starting point for the sampling algorithms using five iterations of MATLAB’s genetic algorithm with a population size of 100, resulting in 600 full FSP evaluations. We then refine the output of the genetic algorithm with a local search using *fmincon* with a maximum of 1000 further evaluations of the full model. This is a negligible cost in comparison to the 10, 000 iterations that we set for the sampling algorithms.

**Table 2:** Bounds on the support of the prior distribution of the parameter in the Two-state gene expression example.

| Parameter | $k_{\text{on}}$ | $k_{\text{off}}$ | $k_r$ | $\gamma$ |
| --- | --- | --- | --- | --- |
| Lower bound | 1.00e-06 | 1.00e-06 | 1.00e-06 | 1.00e-06 |
| Upper bound | 1.00e+01 | 1.00e+01 | 1.00e+04 | 1.00e+01 |

Prior is uniform on the $\log_{10}$ -transformed parameter space. Parameter units are $\text{hr}^{-1}$ .

We summarize the performance characteristics of the sampling schemes in Table 3. The ADAMH-FSP-Krylov requires less computational time (Fig. 1) without a significant reduction in the multivariate effective sample size. In terms of computational time, the ADAMH-FSP-Krylov takes less time to generate an independent sample. This is partly explained by observing that the first stage of the scheme filters out many low-density samples with the efficient approximation, resulting in 81.09% fewer full evaluations in the second stage (cf. Table 3).

**Table 3:** Performance of the sampling algorithms on the Two-state gene expres-sion example.

| | mESS | CPU<br>time<br>(sec) | $\frac{\text{CPU time}}{\text{mESS}}$ (sec) | Number of<br>full evaluations | Number of<br>rejections | Number of<br>rejections by<br>full FSP |
| --- | --- | --- | --- | --- | --- | --- |
| AMH-FSP | 6078.61 | 20319.33 | 3.34 | 100000 | 79152 | 79152 |
| ADAMH-FSP-Krylov | 5141.64 | 11243.99 | 2.19 | 18905 | 81193 | 98 |
| Hybrid | 5104.85 | 8226.57 | 1.61 | 2111 | 79378 | 7 |

Each algorithm was run for $10^5$ iterations.

**Figure 1:**
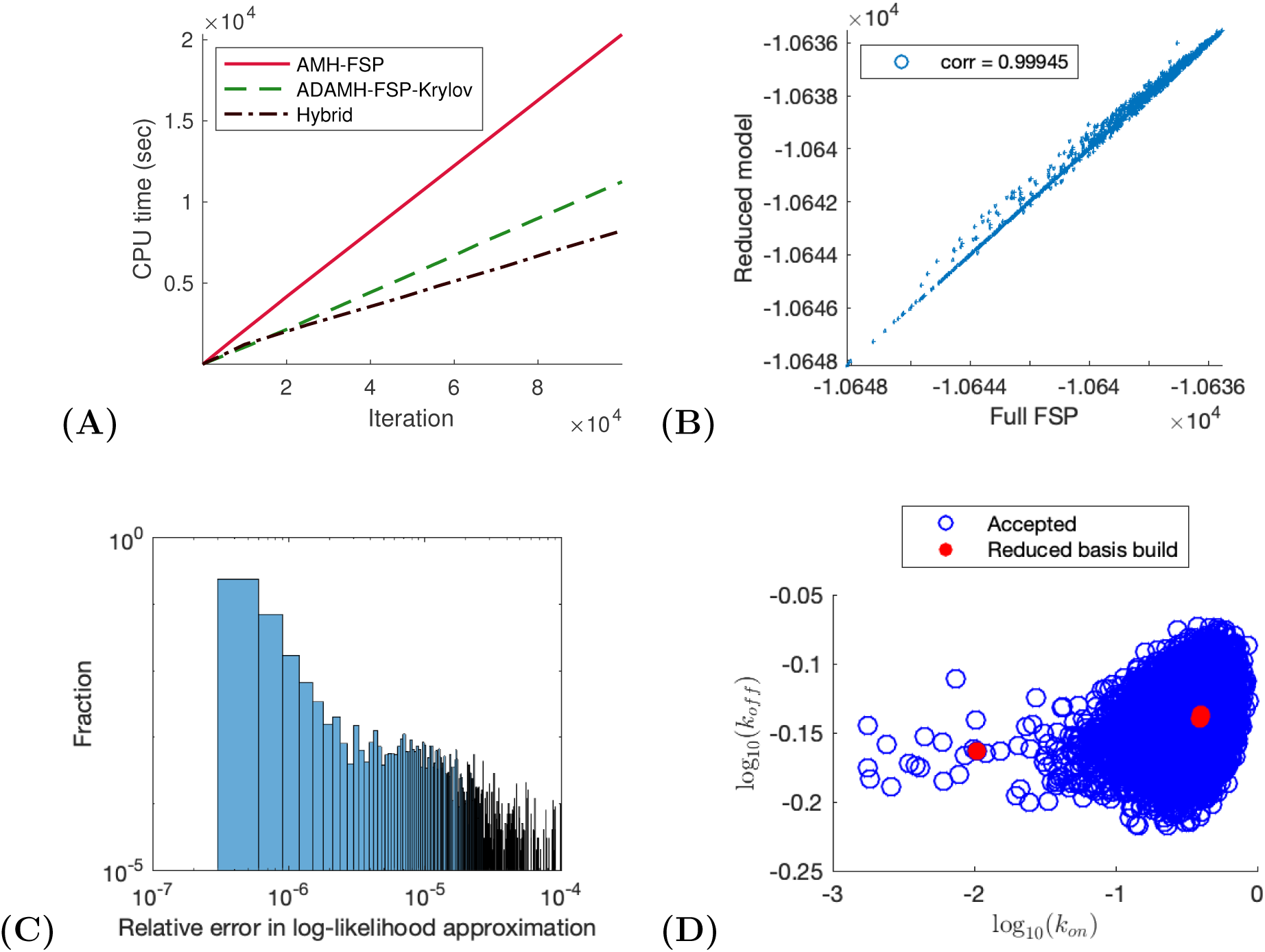
Two-state gene expression example. **(A)** CPU time vs number of iterations for a sample run of the ADAMH-FSP-Krylov and the AMH-FSP. **(B)** Scatterplot of the unno-marlized log-posterior evaluated using the full FSP and the reduced model. Notice that the approximate and true values are almost identical with a correlation coefficient of approximately 0.99853. **(C)** Distribution of the relative error in the approximate log-likelihood evaluations at the parameters accepted by the ADAMH chain. **(D)** 2-D projections of parameter combinations accepted by the ADAMH scheme (blue) and parameter combinations used for reduced model construction (red).

We observe from the scatterplot of log-posterior values of the parameters accepted by the ADAMH-FSP-Krylov that the reduced model evaluations are very close to the FSP evaluations, with the majority of the approximate log-posterior values having a relative error below 10^−4^, with an average of 9.39 *×* 10^−7^ and a median of 2.26 *×* 10^−7^ across all 2152 accepted parameter combinations (Fig. 1 C). This accuracy is achieved with a reduced set of no more than 152 basis vectors per time subinterval that was built using solutions from only *four* sampled parameter combinations (Fig. 2). All the basis updates occur during the first tenth portion of the chain, and these updates consume less than one percent of the total chain runtime (Table 4).

**Table 4:** Breakdown of CPU time spent in the main components of ADAMH-FSP-Krylov run in the Two-state gene expression example.

| Component | Time occupied (sec) | Fraction of total time (per cent) |
| --- | --- | --- |
| Full FSP Evaluation | 3632.34 | 32.30 |
| Reduced Model Evaluation | 7402.34 | 65.83 |
| Reduced Model Update | 5.07 | 0.05 |
| Total | 11243.99 | 100.00 |

**Figure 2:**
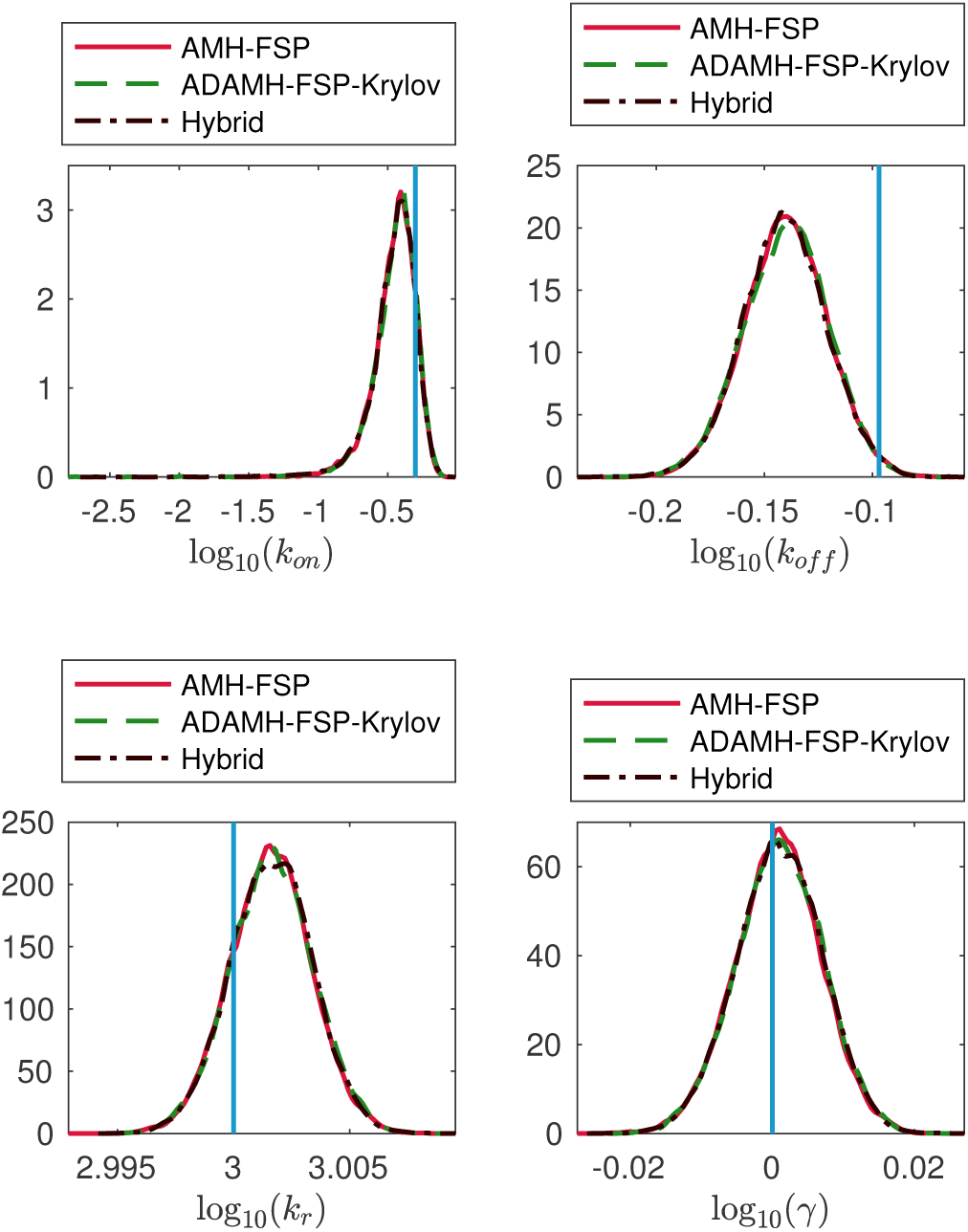
Two-state gene expression example. Estimations of the marginal posterior distributions of the parameters *k*_*on*_, *k*_*off*_, *k*_*r*_, *γ* using the Adaptive Delayed Acceptance Metropolis-Hastings with Krylov reduced model (ADAMH-FSP-Krylov), the Adaptive Metropolis-Hastings with full FSP (AMH-FSP), and the Hybrid method.

**Figure 3:**
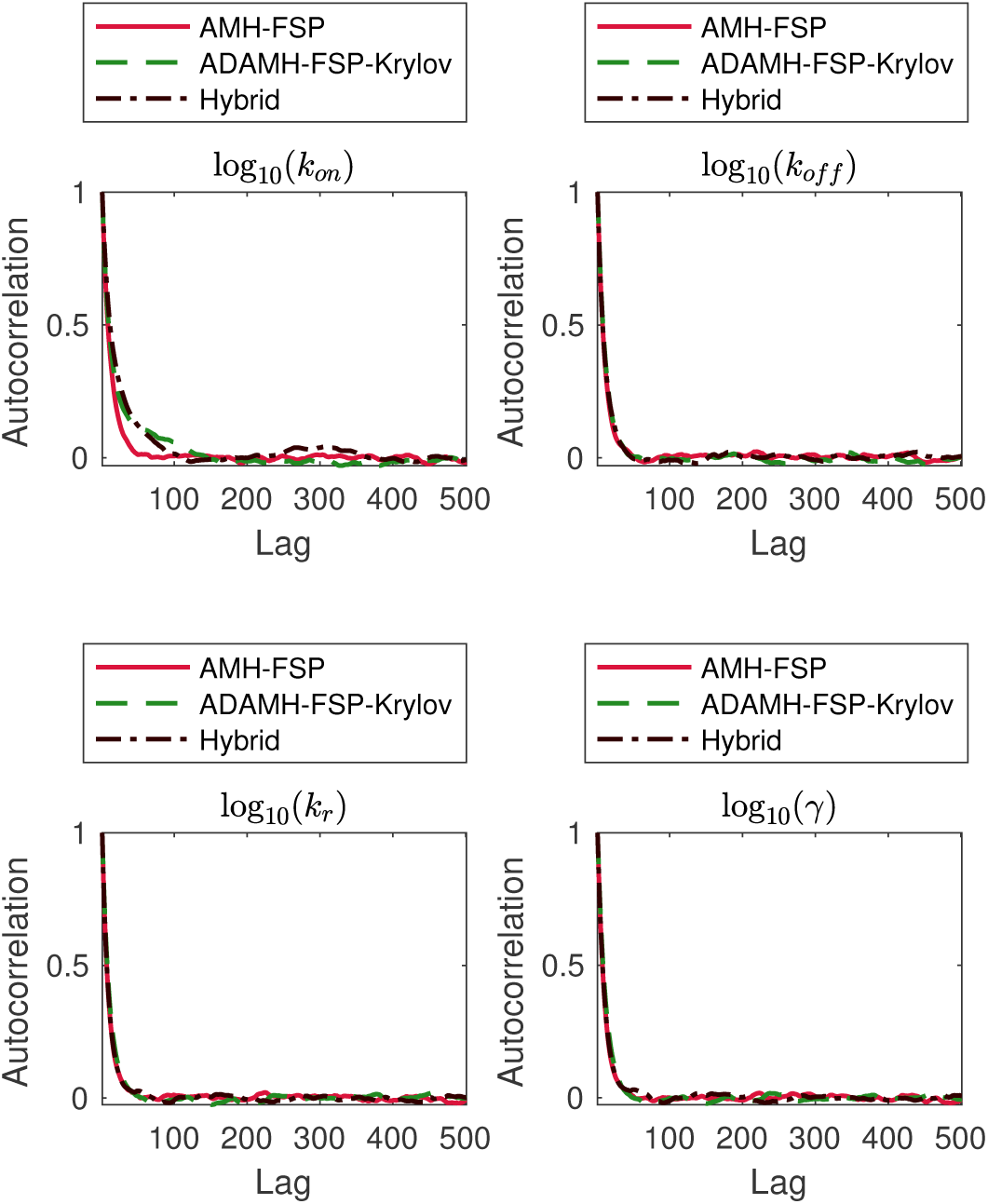
Two-state gene expression example. Autocorrelation in the outputs of the Adaptive Delayed Acceptance Metropolis-Hastings with Krylov reduced model (ADAMH-FSP-Krylov), the Adaptive Metropolis-Hastings with full FSP (AMH-FSP), and the Hybrid method for the parameters *k*_*on*_, *k*_*off*_, *k*_*r*_, *γ*. The autocorrelation functions are estimated in the *log*_10_-transformed space of the parameters, in which the three chains draw their proposals. Actual chain lengths are 10^5^, which is approximately 2500-fold longer than the longest decorrelation time (cf. Table 6).

From the samples obtained by the ADAMH-Krylov-FSP, we found that full and reduced FSP evaluation take approximately 0.19 and 0.07 seconds on average, allowing for a maximal speedup factor of approximately 100(0.19 - 0.07)/0.19 *≈* 61.51% for the current model reduction scheme. Here, the term *reduced model* refers to the final reduced model obtained from the adaptive reduced basis update of the ADAMH-Krylov-FSP. The speedup offered by the ADAMH-Krylov-FSP was found to be 100(20319.33 - 11243.99)/20319.33 *≈* 44.66%, or approximately two thirds the maximal achievable improvement for the current model reduction scheme. Interestingly, the Hybrid scheme yields almost identical results using the reduced model alone in comparison to using the full model (Fig. 2 and Table 5), and the 65.03% reduction in computational effort matched very well to the maximal estimated improvement.

**Table 5:** Posterior mean and standard deviation of the Two-state gene expression example estimated by the sampling schemes.

| Parameter | AMH-FSP |  | ADAMH-FSP-Krylov |  | Hybrid |  | True |
| --- | --- | --- | --- | --- | --- | --- | --- |
|  | mean | std | mean | std | mean | std |  |
| $\log_{10}(k_{\text{on}})$ | -4.52e-01 | 1.50e-01 | -4.51e-01 | 1.66e-01 | -4.58e-01 | 1.71e-01 | -3.01e-01 |
| $\log_{10}(k_{\text{off}})$ | -1.40e-01 | 1.92e-02 | -1.40e-01 | 1.95e-02 | -1.41e-01 | 1.93e-02 | -9.69e-02 |
| $\log_{10}(k_r)$ | 3.00e+00 | 1.76e-03 | 3.00e+00 | 1.78e-03 | 3.00e+00 | 1.76e-03 | 3.00e+00 |
| $\log_{10}(\gamma)$ | 8.33e-04 | 6.05e-03 | 1.03e-03 | 6.09e-03 | 9.85e-04 | 6.06e-03 | 0.00e+00 |

Parameter values are shown in $\log_{10}$ scale.
All parameters have units $\text{hr}^{-1}$ .
The ‘True’ column shows the true parameter values.

**Table 6:** Geweke diagnostics and integrated autocorrelation time for the three chains in the Two-state gene expression example.

| Parameter | AMH-FSP |  | ADAMH-FSP-Krylov |  | Hybrid |  |
| --- | --- | --- | --- | --- | --- | --- |
|  | Geweke | IACT | Geweke | IACT | Geweke | IACT |
| $\log_{10}(k_{\text{on}})$ | 9.92e-01 | 2.32e+01 | 9.25e-01 | 4.10e+01 | 9.90e-01 | 4.01e+01 |
| $\log_{10}(k_{\text{off}})$ | 9.97e-01 | 1.98e+01 | 9.99e-01 | 2.11e+01 | 9.92e-01 | 2.04e+01 |
| $\log_{10}(k_r)$ | 1.00e+00 | 1.86e+01 | 1.00e+00 | 2.04e+01 | 1.00e+00 | 1.93e+01 |
| $\log_{10}(\gamma)$ | 7.55e-01 | 1.93e+01 | 6.87e-01 | 2.10e+01 | 7.01e-01 | 2.07e+01 |

Geweke column shows the p-values in the Geweke diagnostics. This test is passed at 5% significance level if the p-value is above 0.05.
IACT column shows the integrated autocorrelation time, which is the number of samples in the MCMC chain that are equivalent to an independent posterior sample.

### Genetic toggle switch

The next model we consider in our numerical tests is the nonlinear genetic toggle switch, which is based on a well-known synthetic gene circuit that consists of two genes whose products repress each other.^63^ In the model that we consider, ^64,65^ the gene products are represented by two species where each species’ production rate is repressed by the copy number of the other species according to a Hill function (Table 7). As a result when one gene has high activity, the other is repressed and vice versa. Using the stochastic simulations and the ‘true’ parameters as given in Table 8 (we use the same parameters as those in Fox and Munsky^66^), we generate data at 2, 6 and 8 hours, each with 500 single-cell samples. To build the reduced bases for the FSP reduction, we use the union of ten equally-spaced points between zero and 8 hrs and the time points of observations. The prior distribution in our test was chosen as the log-uniform distribution on a rectangle, whose bounds are given in Table 9. The full FSP size is set as the rectangle *{*0, *…*, 100*} × {*0, *…*, 100*}*, corresponding to 10,201 states.

**Table 7:** Genetic toggle switch reactions and propensities.

|  | reaction | propensity |
| --- | --- | --- |
| 1. | $X \longrightarrow \emptyset$ | $\gamma_X[X]$ |
| 2. | $\emptyset \longrightarrow X$ | $k_{0X} + \frac{k_{1X}}{1+a_{yx}([Y]^{n_{yx}})}$ |
| 3. | $Y \longrightarrow \emptyset$ | $\gamma_Y[Y]$ |
| 4. | $\emptyset \longrightarrow Y$ | $k_{0Y} + \frac{k_{1Y}}{1+a_{xy}([X]^{n_{xy}})}$ |

The units for the parameters $k_{0X}, k_{1X}, \gamma_X, k_{0Y}, k_{1Y}, \gamma_Y$ are $\text{sec}^{-1}$ . The other parameters (dimensionless) are fixed at $a_{yx} = 2.6 \times 10^{-3}, a_{xy} = 6.1 \times 10^{-3}, n_{yx} = 3, n_{xy} = 2.1$

**Table 8:** Posterior mean and standard deviation of the Genetic toggle switch parameters estimated by the sampling schemes.

| Parameter | AMH-FSP |  | ADAMH-FSP-Krylov |  | Hybrid |  | True |
| --- | --- | --- | --- | --- | --- | --- | --- |
|  | mean | std | mean | std | mean | std |  |
| $\log_{10}(k_{0X})$ | -2.65e+00 | 2.79e-02 | -2.65e+00 | 2.83e-02 | -2.65e+00 | 2.79e-02 | -2.66e+00 |
| $\log_{10}(k_{1X})$ | -1.75e+00 | 2.27e-02 | -1.75e+00 | 2.31e-02 | -1.75e+00 | 2.23e-02 | -1.77e+00 |
| $\log_{10}(\gamma_X)$ | -3.40e+00 | 2.53e-02 | -3.40e+00 | 2.57e-02 | -3.40e+00 | 2.51e-02 | -3.42e+00 |
| $\log_{10}(k_{0Y})$ | -5.05e+00 | 5.45e-01 | -5.05e+00 | 5.44e-01 | -5.08e+00 | 5.40e-01 | -4.17e+00 |
| $\log_{10}(k_{1Y})$ | -1.77e+00 | 2.14e-02 | -1.77e+00 | 2.19e-02 | -1.77e+00 | 2.13e-02 | -1.80e+00 |
| $\log_{10}(\gamma_Y)$ | -3.39e+00 | 2.26e-02 | -3.39e+00 | 2.31e-02 | -3.39e+00 | 2.25e-02 | -3.42e+00 |

True parameter values are shown in the ‘True’ column.
Parameter values are shown in $\log_{10}$ scale.
All parameters shown have unit $\text{sec}^{-1}$ .

**Table 9:** Bounds on the support of the prior in the Genetic toggle switch example.

| Parameter | $k_{0X}$ | $k_{1X}$ | $\gamma_X$ | $k_{0Y}$ | $k_{1Y}$ | $\gamma_Y$ |
| --- | --- | --- | --- | --- | --- | --- |
| Lower bound | 1.00e-06 | 1.00e-06 | 1.00e-06 | 1.00e-06 | 1.00e-06 | 1.00e-06 |
| Upper bound | 1.00e-01 | 1.00e-01 | 1.00e-01 | 1.00e-01 | 1.00e-01 | 1.00e-01 |

Parameters have the same unit $\text{sec}^{-1}$ .
Prior is uniform in the $\log_{10}$ -transformed parameter space, covering five orders of magnitude for each parameter.

To find the starting point for the chains, we run five generations of MATLAB’s genetic algorithm with 1100 full FSP evaluations. Then, we run another 600 iterations of *fmincon* to refine the output of the *ga* solver. Using the parameter vector output by this combined optimization scheme as initial sample, we run both the ADAMH-FSP-Krylov and the AMH-FSP for 100, 000 iterations.

From the samples obtained by the ADAMH, we found that Expokit takes 0.29 sec to solve the full FSP model and 0.08 sec to solve the reduced model. This results in a maximal potential savings of about 71.78% when exclusively using the reduced FSP model.

The efficiency of the ADAMH-Krylov-FSP is confirmed in Table 10, where the delayed acceptance scheme is 49.85% faster than the AMH-FSP algorithm. Similar to the last example, we observe a close agreement between the first and second stage of the ADAMH run, where 96.41% of the proposals promoted by the reduced-model-based evaluations are accepted by the full-FSP-based evaluation. This high second-stage acceptance rate is explained by the quality of the reduced model in approximating the log-posterior values (Fig. 4 C). The accurate reduced model, constructed from only *ten* parameter samples, consists of no more than 530 basis vectors per time subinterval, with all the basis updates occurring during the first tenth portion of the chain.

**Table 10:** Performance of the sampling algorithms applied to the Genetic toggle switch example.

| | mESS | CPU time (sec) | $\frac{\text{CPU time}}{\text{mESS}}$ (sec) | Number of full evaluations | Number of rejections | Number of rejections by full FSP |
| --- | --- | --- | --- | --- | --- | --- |
| AMH-FSP | 4343.31 | 30238.78 | 6.96 | 100000 | 76267 | 76267 |
| ADAMH-FSP-Krylov | 4299.54 | 15165.87 | 3.53 | 23530 | 77314 | 844 |
| Hybrid | 4276.67 | 7100.20 | 1.66 | 2915 | 76169 | 276 |

The total chain length for each algorithm was $10^5$ .
The ADAMH-FSP-Krylov scheme uses markedly fewer full evaluations than the AMH-FSP scheme, and 96.41% of the parameters promoted by the first-stage are accepted in the second stage.

**Figure 4:**
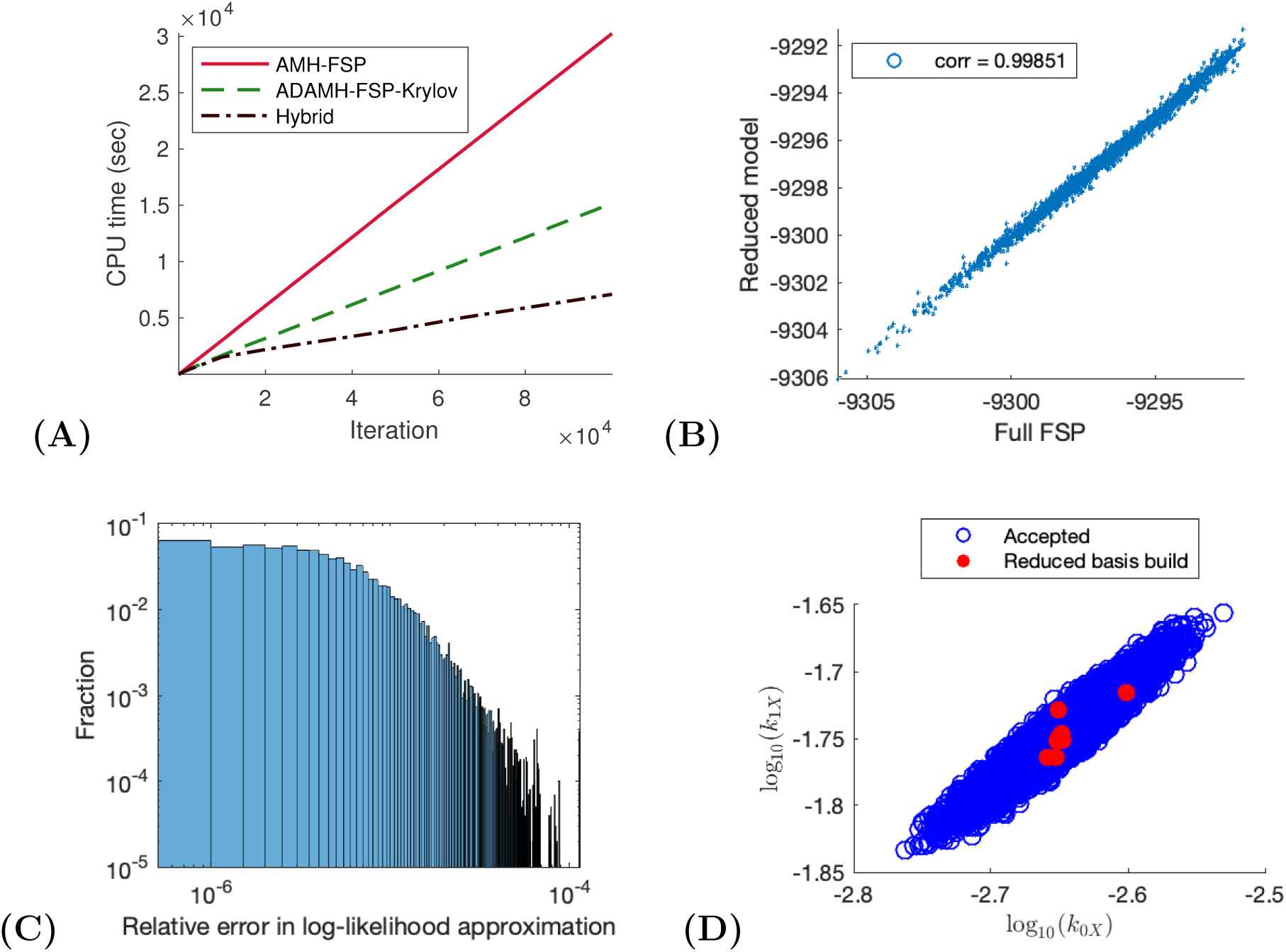
Genetic toggle switch example. (**A**) CPU time vs number of iterations for a sample run of the ADAMH-FSP-Krylov and the AMH-FSP. (**B**) Scatterplot of the unnomarlized log-posterior evaluated using the full FSP and the reduced model. Notice that the approximate and true values are almost identical with a correlation coefficient of approximately 0.99851. (**C**) Distribution of the relative error in the approximate log-likelihood evaluations at the parameters accepted by the ADAMH chain. (**D**) 2-D projections of parameter combinations accepted by the ADAMH scheme (blue) and parameter combinations used for reduced model construction (red).

The effectiveness of the ADAMH’s model-building procedure explains the good behavior of the Hybrid algorithm, which yields similar results to the reference chain (Fig. 5) but with reduced computational time (Table 10). The Hybrid scheme achieves a saving of 60.50%, which is close to the estimated maximum speedup of 71.78% mentioned above.

**Figure 5:**
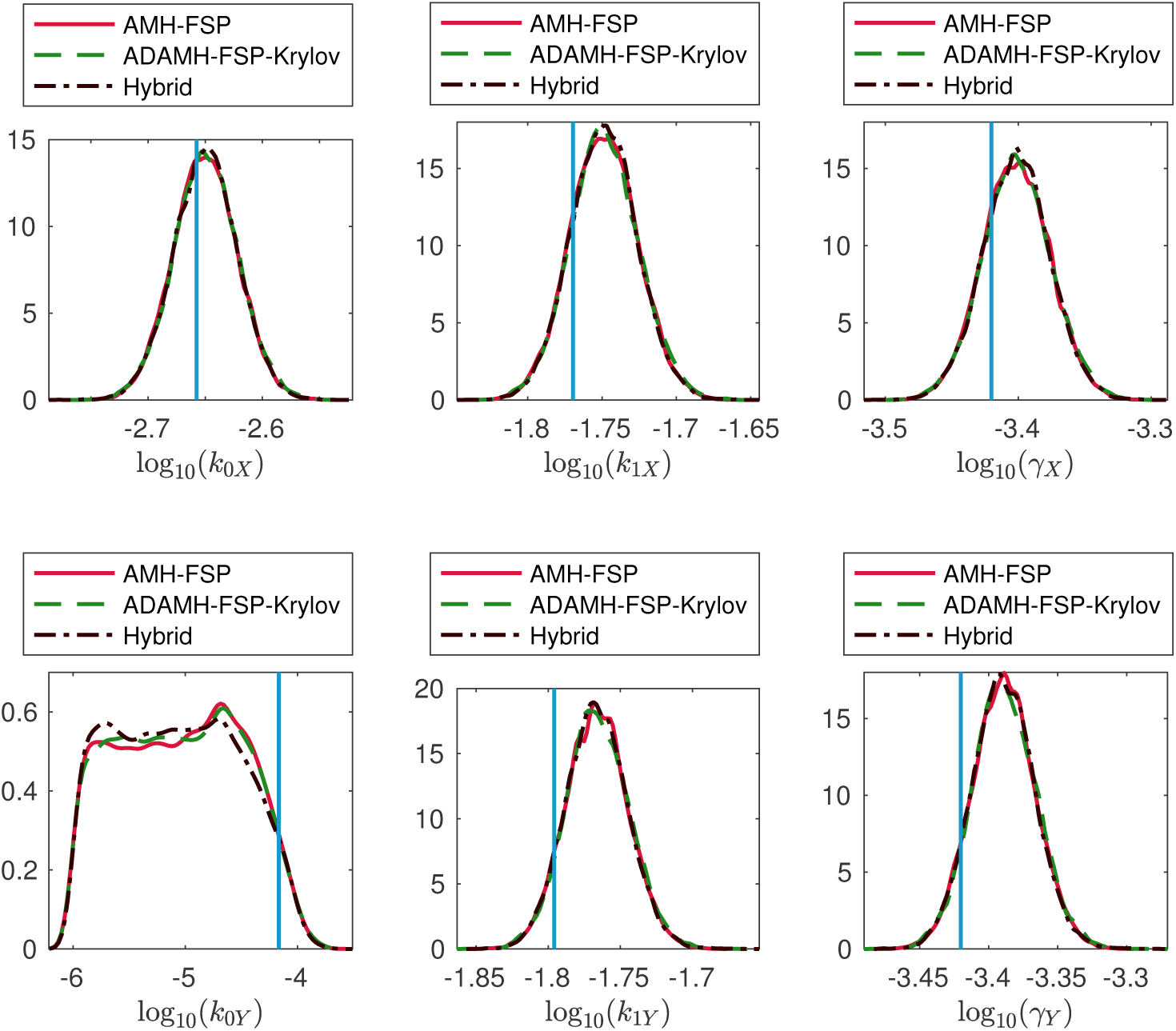
Genetic toggle switch example. Estimations of the marginal posterior distributions of the parameters *k*_0*X*_, *k*_1*X*_, *γ*_*X*_, *k*_0*Y*_, *k*_1*Y*_, *γ*_*Y*_ using the Adaptive Delayed Acceptance Metropolis-Hastings with Krylov reduced model (ADAMH-FSP-Krylov), the Adaptive Metropolis-Hastings with full FSP (AMH-FSP), and the Hybrid method.

**Figure 6:**
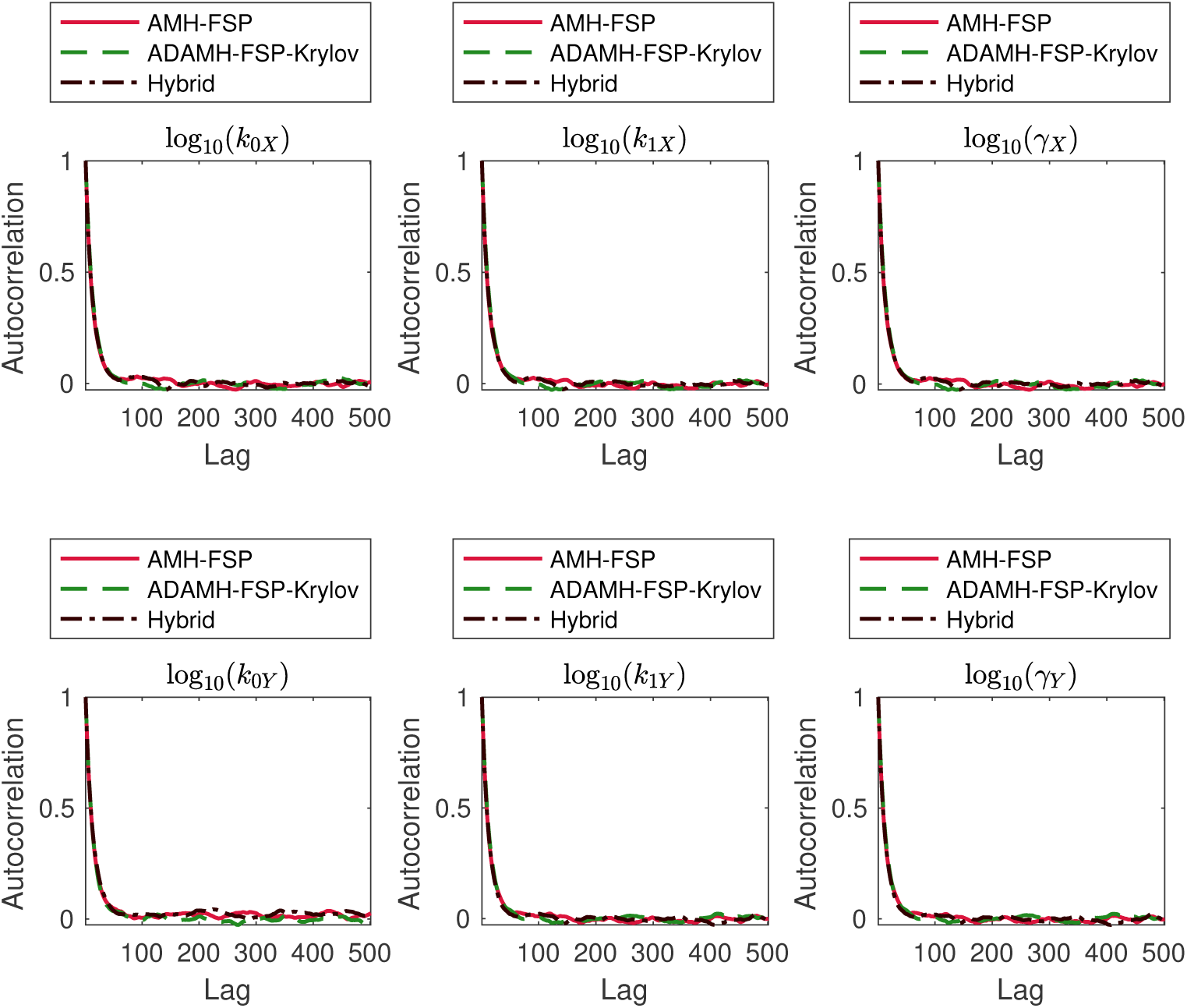
Genetic toggle switch example. Autocorrelation in the outputs of the Adaptive Delayed Acceptance Metropolis-Hastings with Krylov reduced model (ADAMH-FSP-Krylov), the Adaptive Metropolis-Hastings with full FSP (AMH-FSP), and the Hybrid method for the parameters *k*_0*X*_, *k*_1*X*_, *γ*_*X*_, *k*_0*Y*_, *k*_1*Y*_, *γ*_*Y*_. The autocorrelation is estimated directly in log_10_-transformed space of parameters, where the chains draw their proposals.

### A gene expression model with spatial components

The last two demonstrative examples allow us to vaildate the accuracy and to some extent the computatial speedups brought by the new approximate sampling schemes. We now consider an example where the exact Bayesian analysis is no longer practical. We extend the gene expression model described above to distinguish between the nucleus and cytoplasmic compartments in the cell, similar to a stochastic model recently considered for MAPK-activated gene expression dynamics in yeast. ^15^ The gene can transition between four states {0, 1, 2, 3} with transcription activated when the gene state is in states 1 to 3. RNA is transcribed in the nucleus and later transported to the cytoplasm as a first order reaction. These cellular processes and the degradation of RNA in both spatial compartments are modeled by a reaction network with six reactions and three species (Table 13).

**Table 11:** Breakdown of CPU time spent in the main components of ADAMH-FSP-Krylov run in the Genetic toggle switch example.

| Component | Time occupied (sec) | Fraction of total time (per cent) |
| --- | --- | --- |
| Full FSP Evaluation | 6774.00 | 44.67 |
| Reduced Model Evaluation | 8121.47 | 53.55 |
| Reduced Model Update | 23.39 | 0.15 |
| Total | 15165.87 | 100.00 |

**Table 12:** Convergence diagnostics for the three sampling algorithms applied to the Genetic toggle switch example.

| Parameter | AMH-FSP |  | ADAMH-FSP-Krylov |  | Hybrid |  |
| --- | --- | --- | --- | --- | --- | --- |
|  | Geweke | IACT | Geweke | IACT | Geweke | IACT |
| $\log_{10}(k_{0X})$ | 1.00e+00 | 2.57e+01 | 1.00e+00 | 2.68e+01 | 1.00e+00 | 2.61e+01 |
| $\log_{10}(k_{1X})$ | 1.00e+00 | 2.59e+01 | 1.00e+00 | 2.72e+01 | 1.00e+00 | 2.54e+01 |
| $\log_{10}(\gamma_X)$ | 1.00e+00 | 2.56e+01 | 1.00e+00 | 2.69e+01 | 1.00e+00 | 2.60e+01 |
| $\log_{10}(k_{0Y})$ | 9.78e-01 | 2.79e+01 | 9.91e-01 | 2.70e+01 | 9.93e-01 | 2.85e+01 |
| $\log_{10}(k_{1Y})$ | 1.00e+00 | 2.65e+01 | 1.00e+00 | 2.64e+01 | 9.99e-01 | 2.47e+01 |
| $\log_{10}(\gamma_Y)$ | 1.00e+00 | 2.64e+01 | 1.00e+00 | 2.62e+01 | 1.00e+00 | 2.46e+01 |

Geweke column shows the p-values in the Geweke diagnostics. This test is passed at 5% significance level if the p-value is above 0.05.
IACT column shows the integrated autocorrelation time, which is the number of samples in the MCMC chain that are equivalent to an independent posterior sample.

**Table 13:** Spatial gene expression reactions and propensities.

| reaction | propensity |
| --- | --- |
| 1. $G_i \xrightarrow{k_{gene}^+} G_{i+1}$ | $\alpha_1 = k_{gene}^+[i \leq 2]$ |
| 2. $G_i \xrightarrow{k_{gene}^-} G_{i-1}$ | $\alpha_2 = k_{gene}^-[i \geq 1]$ |
| 3. $G_i \xrightarrow{k_r} G_i + RNA_{nuc}$ | $\alpha_3 = k_r[i \geq 1]$ |
| 4. $RNA_{nuc} \xrightarrow{\gamma_{nuc}} \emptyset$ | $\alpha_4 = \gamma_{nuc}[RNA_{nuc}]$ |
| 5. $RNA_{nuc} \xrightarrow{k_{trans}} RNA_{cyt}$ | $\alpha_5 = k_{trans}[RNA_{nuc}]$ |
| 6. $RNA_{cyt} \xrightarrow{\gamma_{cyt}} \emptyset$ | $\alpha_6 = \gamma_{cyt}[RNA_{cyt}]$ |

The gene is considered as one species with 4 different states $G_i$ , $i = 0, \dots, 3$ . Parameters' units are $\text{sec}^{-1}$ .

We simulated a data set of 100 single-cell measurements at five equally-spaced time points between one and 10 minute(s) (min), that is, *T*_*data*_ = {2, 4, 6, 8, 10} (min). The time points for generating the basis are *T*_*basis*_ = *T*_*data*_ *∪* {*j ×* 0.2 min, *j* = 1, *…*, 50}. We chose the basis update threshold as *δ* = 10^−4^. The prior distribution in our test is the log-uniform distribution on a rectangle, whose bounds are given in Table 14. The full FSP state space is chosen as

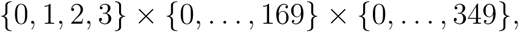

which results in 238, 000 states. We only run the ADAMH-FSP-Krylov and the Hybrid schemes, starting from the reference parameters.

**Table 14:** Bounds on the support of the prior in the Spatial gene expression example.

| Parameter | $k_{\text{gene}}^+$ | $k_{\text{gene}}^-$ | $k_r$ | $\gamma_{\text{nuc}}$ | $k_{\text{trans}}$ | $\gamma_{\text{cyt}}$ |
| --- | --- | --- | --- | --- | --- | --- |
| Lower bound | 1.00e-06 | 1.00e-06 | 1.00e-06 | 1.00e-08 | 1.00e-06 | 1.00e-06 |
| Upper bound | 1.00e+00 | 1.00e+00 | 1.00e+01 | 1.00e+00 | 1.00e+00 | 1.00e+00 |

All parameters have the same unit $\text{sec}^{-1}$ .
Prior is uniform in the $\log_{10}$ -transformed parameter space.

Inspecting the autocorrelation plots (Fig. 9) and performing Geweke diagnostics on the two chain outputs (Table 18) did not reveal any convergence issues. Table 15 summarizes the performance of the two algorithms for 10^4^ iterations. From the posterior samples of the ADAMH chain, we estimate that an average full FSP evaluation would take 101.82 seconds. Therefore, we estimate that running the AMH-FSP to the same number of iterations as the other two chains would have taken more than one week to finish (about 282 hours *≈* days). In contrast, the ADAMH-FSP-Krylov scheme took about 93.9 hours (about four days) to finish, resulting in a reduction of 66.7%, or a speedup factor of three, in terms of total computational time.

**Table 15:** Performance of the sampling algorithms applied to the Spatial gene expression example.

| | mESS | CPU<br>time<br>(sec) | $\frac{\text{CPU time}}{\text{mESS}}$ (sec) | Number of<br>full evaluations | Number of<br>rejections | Number of<br>rejections by<br>full FSP |
| --- | --- | --- | --- | --- | --- | --- |
| ADAMH-FSP-Krylov | 455.96 | 337955.28 | 741.20 | 2863 | 7408 | 271 |
| Hybrid | 470.37 | 92990.58 | 197.70 | 407 | 7192 | 46 |

The total chain length for each algorithm is $10^5$ .

For the ADAMH-FSP-Krylov chain, the log-posterior evaluations from the reduced model are accurate (Fig. 7 C and Table 16), with relative error below the algorithmic tolerance of 10^−4^, with a mean of 4.09 *×* 10^−5^ and a median of 2.45 *×* 10^−5^. This accurate model was built automatically by the ADAMH scheme using just 17 points in the parameter space (Fig. 7 D), resulting in a set of no more than 538 vectors per time subinterval. All the basis updates occur during the first fifth portion of the chain, and these updates consume about 5.91% of the total runtime (Table 17). The high accuracy of the posterior approximation translates into a very high second-stage acceptance of 90.53% of the proposals promoted by the first-stage reduced-model-based evaluation. Such high acceptance rates in the second stage are crucial to the efficiency for the delayed acceptance scheme, since almost all of the expensive FSP evaluations are accepted. ^35^

**Table 16:** Posterior mean and standard deviation of the Spatial gene expression parameters estimated by the sampling schemes.

| Parameter | ADAMH-FSP-Krylov |  | Hybrid |  | True |
| --- | --- | --- | --- | --- | --- |
|  | mean | std | mean | std |  |
| $\log_{10}(k_{\text{gene}}^+)$ | -2.51e+00 | 3.73e-02 | -2.52e+00 | 3.82e-02 | -2.52e+00 |
| $\log_{10}(k_{\text{gene}}^-)$ | -2.18e+00 | 2.88e-02 | -2.19e+00 | 2.91e-02 | -2.22e+00 |
| $\log_{10}(k_r)$ | 1.71e-01 | 8.98e-03 | 1.70e-01 | 9.80e-03 | 1.76e-01 |
| $\log_{10}(\gamma_{\text{nuc}})$ | -2.55e+00 | 6.42e-02 | -2.56e+00 | 6.98e-02 | -2.52e+00 |
| $\log_{10}(k_{\text{trans}})$ | -2.00e+00 | 8.23e-03 | -2.00e+00 | 8.49e-03 | -2.00e+00 |
| $\log_{10}(\gamma_{\text{cyt}})$ | -2.54e+00 | 1.72e-02 | -2.54e+00 | 1.84e-02 | -2.52e+00 |

Parameter values are shown in $\log_{10}$ scale. All parameters have units $\text{sec}^{-1}$ .
True values are shown in the ‘True’ column.

**Table 17:**
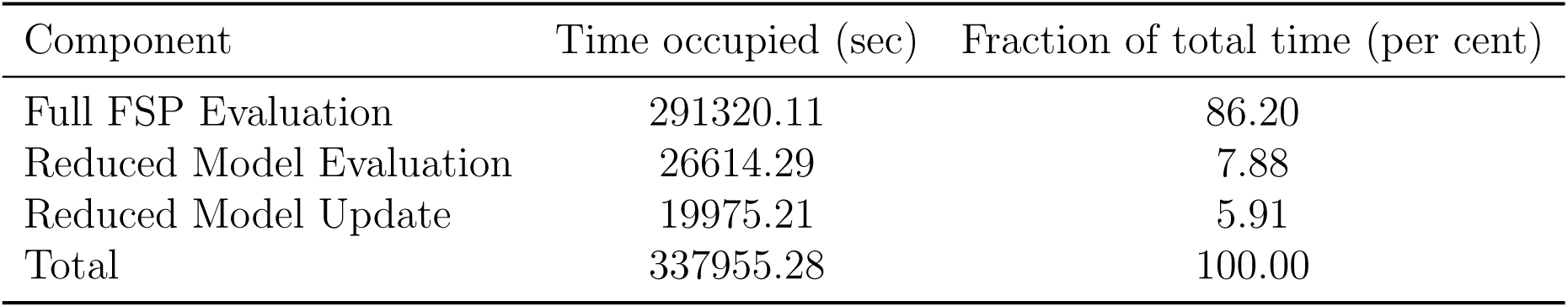
Breakdown of CPU time spent in the maiin components of ADAMH-FSP-Krylov in the Spatial gene expression example.

| Component | Time occupied (sec) | Fraction of total time (per cent) |
| --- | --- | --- |
| Full FSP Evaluation | 291320.11 | 86.20 |
| Reduced Model Evaluation | 26614.29 | 7.88 |
| Reduced Model Update | 19975.21 | 5.91 |
| Total | 337955.28 | 100.00 |

**Table 18:** Convergence diagnostics for the three sampling algorithms applied to the Spatial gene expression example.

| Parameter | ADAMH-FSP-Krylov |  | Hybrid |  |
| --- | --- | --- | --- | --- |
|  | Geweke | IACT | Geweke | IACT |
| $\log_{10}(k_{\text{gene}}^+)$ | 9.91e-01 | 3.05e+01 | 9.95e-01 | 3.17e+01 |
| $\log_{10}(k_{\text{gene}}^-)$ | 9.90e-01 | 2.78e+01 | 9.92e-01 | 3.87e+01 |
| $\log_{10}(k_r)$ | 9.78e-01 | 2.77e+01 | 9.55e-01 | 3.49e+01 |
| $\log_{10}(\gamma_{\text{nuc}})$ | 9.90e-01 | 3.19e+01 | 9.78e-01 | 3.62e+01 |
| $\log_{10}(k_{\text{trans}})$ | 9.99e-01 | 3.63e+01 | 9.99e-01 | 1.74e+01 |
| $\log_{10}(\gamma_{\text{cyt}})$ | 9.99e-01 | 2.65e+01 | 1.00e+00 | 2.34e+01 |

Geweke column shows the p-values in the Geweke diagnostics. This test is passed at 5% significance level if the p-value is above 0.05.
IACT column shows the integrated autocorrelation time, which is the number of samples in the MCMC chain that are equivalent to an independent posterior sample.

**Figure 7:**
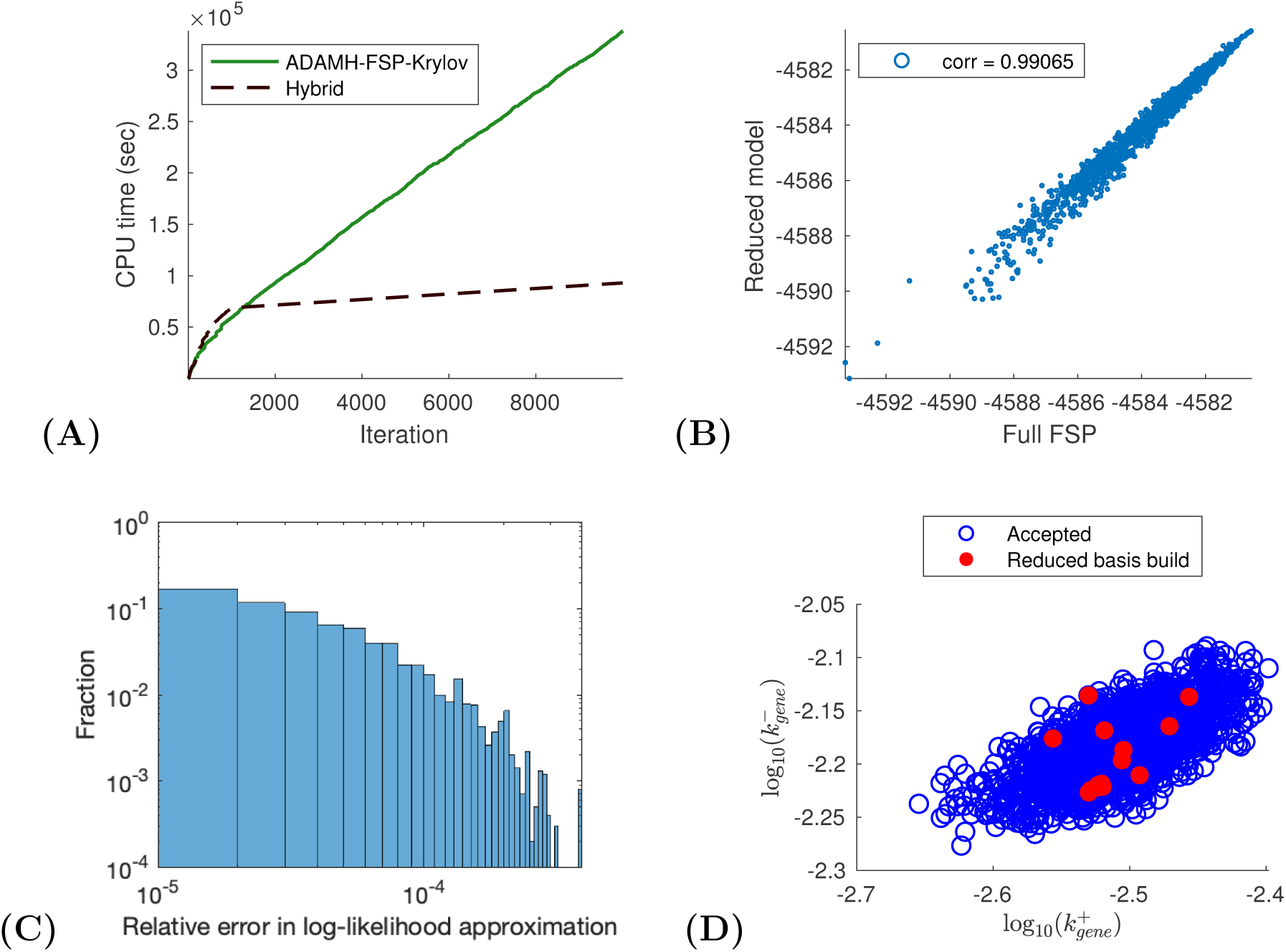
Spatial gene expression example. (**A**) CPU time vs number of iterations for a sample run of the ADAMH-FSP-Krylov and the AMH-FSP. (**B**) Scatterplot of the unno-marlized log-posterior evaluated using the full FSP and the reduced model. Notice that the approximate and true values are almost identical with a correlation coefficient of approximately 0.99851. (**C**) Distribution of the relative error in the approximate log-likelihood evaluations at the parameters accepted by the ADAMH chain. (**D**) 2-D projections of parameter combinations accepted by the ADAMH scheme (blue) and parameter combinations used for reduced model construction (red).

The hybrid scheme took only 26 hours to finish, yielding an estimated reduction of 90% in computational time, that is, an order of magnitude speedup. We note that an average reduced model evaluation takes only 2.66 seconds, leading to a maximum reduction (in terms of total CPU time) of approximately 97.39%, or a speedup factor of 37. The less than ideal speedup in our run is due to the fact that the Hybrid scheme devoted its first 1000 iterations to learn the reduced model. However, as the number of iterations increase, the cost of the learning phase becomes less significant and we expect that the Hybrid scheme would become much more advantageous. Most importantly, switching solely to the reduced model does not incur a significant difference in the parameter estimation results (Table 16 and Fig. 8).

**Figure 8:**
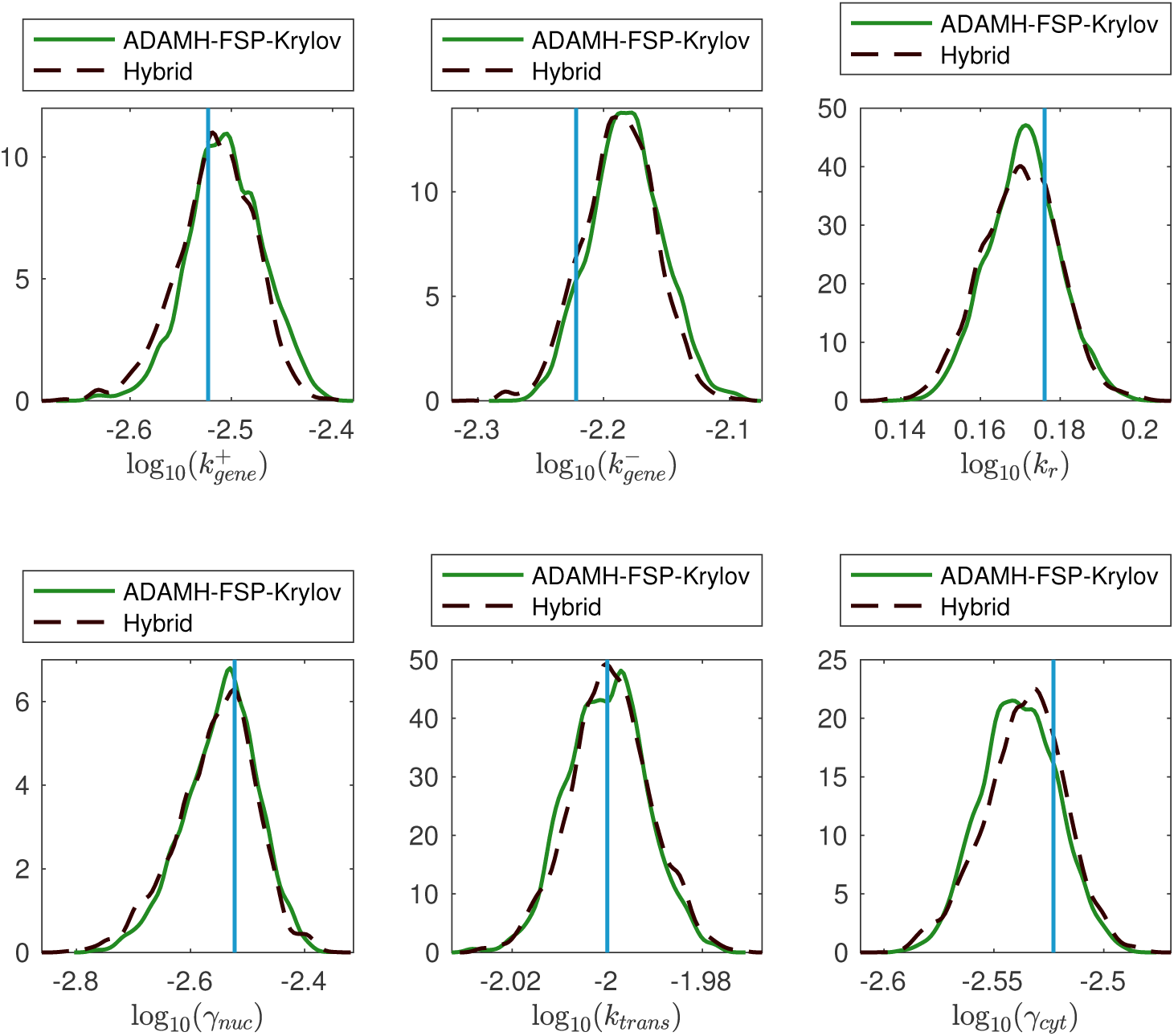
Spatial gene expression example. Estimations of the marginal posterior distributions using the Adaptive Delayed Acceptance Metropolis-Hastings with Krylov reduced model (ADAMH-FSP-Krylov) and the Hybrid method.

**Figure 9:**
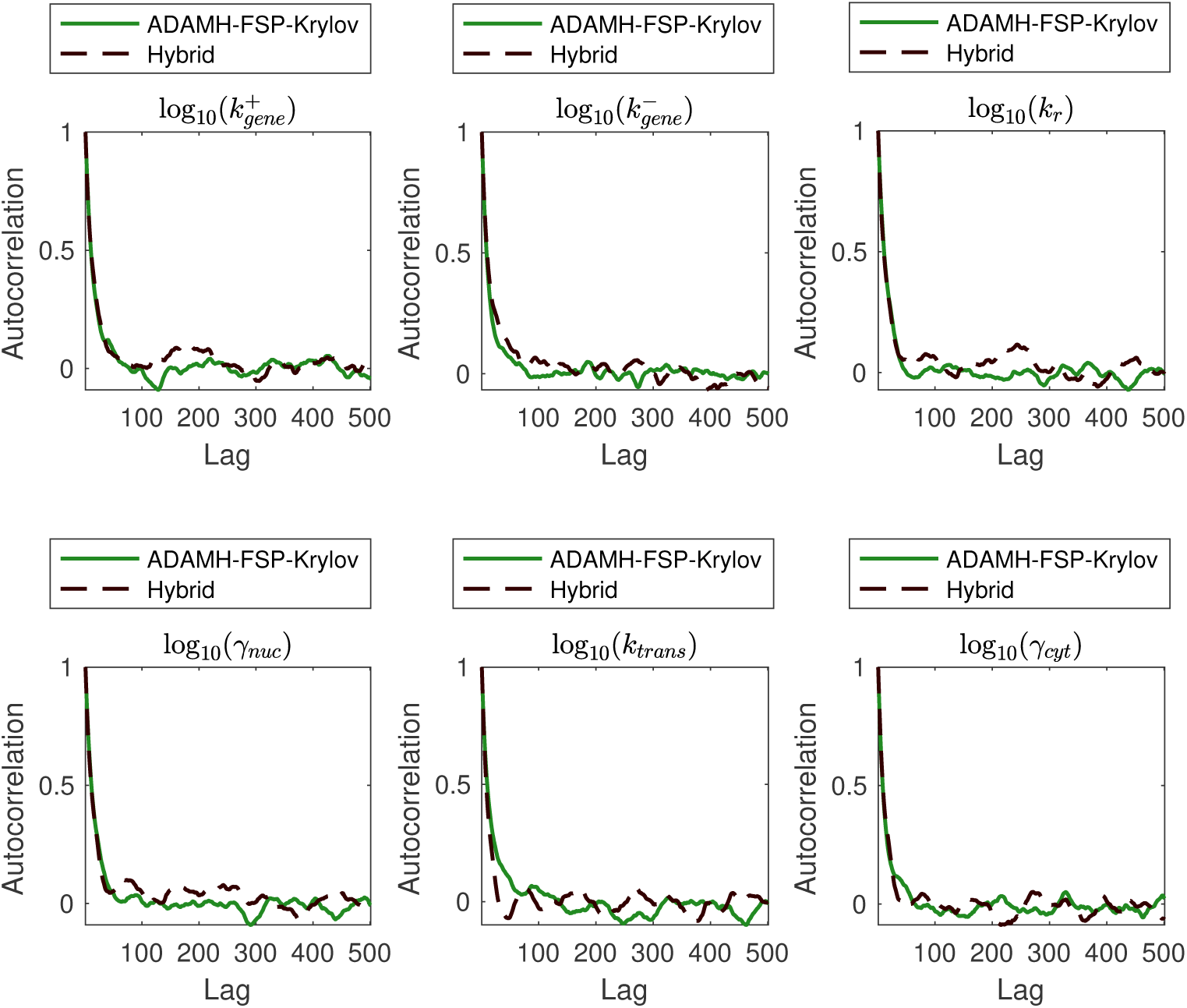
Spatial gene expression example. Autocorrelation in the outputs of the Adaptive Delayed Acceptance Metropolis-Hastings with Krylov reduced model (ADAMH-FSP-Krylov), the Adaptive Metropolis-Hastings with full FSP (AMH-FSP), and the Hybrid method for all six free parameters of the model. The autocorrelation is estimated directly in log_10_-transformed space of parameters, where the chains draw their proposals.

## DISCUSSION

There exists a growing literature on Bayesian inference for temporally resolved stochastic gene expression. These methods concern mainly three types of data analysis: time-course trajectories for individual cells, temporal trajectories of statistical moments, and temporal trajectories of entire populations. Because different types of experimental data provide different kinds of information, they require a diversity of Bayesian methods.

Fluorescence time-course data has attracted the largest interest in stochastic model inference. These data measurements consist of one or more single-cell trajectories observed at discrete time points. The likelihood of the time-course data is usually not directly evaluated but instead estimated via simulations.^37,67–69^ Since the classical stochastic simulation algorithm (SSA)^58^ may be inefficient for large systems with many timescales, many approximation methods have been used to speed up the inference process (see Schnoerr et al.^5^ for an extensive tutorial review on approximation methods for the CME and SSA). Approximate Bayesian Computation (ABC) approaches have also been proposed for time-course data.^70^ We notice that in principle the finite state projection could also be used to directly approximate the log-likelihood for time-course data, and we refer to Andreychenko et al.^71^ for an example that uses an FSP-based likelihood in a frequentist setting. Golightly et al. ^37^ also used a delayed acceptance scheme to analyze time-course data. Their method uses a particle filtering scheme to estimate the first-stage approximate likelihood via the Chemical Langevin equation (CLE) or the Linear Noise approximation (LNA) and the second-stage exact likelihood via the SSA.^58^

Flow cytometry methods collect large amounts of single-cell population data at specific time instances (i.e., snapshots), but lose the temporal correlation of individual cell trajectories. Inference methods based on moments are particularly well suited to these high-throughput experiments thanks to the application of the central limit theorem. ^72,73^ The related methods of Approximate Bayesian Computation (ABC) can also be well utilized for such data.^74^ Recently, Zechner et al.^75^ introduced a hierarchical Bayesian inference framework based on moments of distributions that can incorporate extrinsic noise. A notable challenge in analyzing flow cytometry data is the uncertainty in converting fluorenscence signal into discrete molecular counts. Tiberi et al.^48^ recently introduced a hierarchical Bayesian scheme that enables the inference of measurement noise parameters in addition to model parameters.

Single-molecule optical microscopy approaches such as smFISH provide more precise estimates of molecular counts, but at much lower throughput. smFISH experimens provide spatial, single-molecule resolution, but yields fewer observations than flow cytometry, and in this case, the data are insufficient to accurately estimate the true moments. These moment Single-molecule optical microscopy approaches such as smFISH provide more precise estimates of molecular counts, but at much lower throughput. smFISH experimens provide spatial, single-molecule resolution, but yields fewer observations than flow cytometry, and in this case, the data are insufficient to accurately estimate the true moments. These moment estimation errors can lead to a breakdown of moment-based inference as discussed recently by Munsky et al. ^15^ Maximum likelihood fitting via FSP was found to be able to make much better use of the full information available in these data to effectively constrain the parameters and yield accurate predictions. Unfortunately, full FSP solutions are computationally expensive, which has prevented their wide spread applications. Research on Bayesian and frequentist approaches for analyzing low-count single-cell data appears to be relatively unexplored in comparison to the techniques mentioned above. A few recent examples include the work of Gomez-Schiavon et al. ^10^ and Fox and Munsky^66^ are two recent works that represent respectively a Bayesian and frequentist approach to the analysis and design of smFISH experiments. In this work, we show how MCMC sampling of the posterior from smFISH data can be done efficiently by using reduced order modeling in the delayed acceptance framework.

A surprising observation from our numerical results is that once trained, the reduced model constructed by the ADAMH-FSP-Krylov closely matches the original FSP sampling. This suggests that the ADAMH-FSP-Krylov algorithm could be used as a data-driven method to learn reduced representations of the full FSP-based model, which could then be successfully substituted for the full FSP model in subsequent Bayesian updates. In other words, it could be equally accurate but more efficient to cease full FSP evaluations in the ADAMH scheme once we are confident about the accuracy of the reduced model. In our numerical tests, the ADAMH updates completed within the first 10-20% of the MCMC chain, at which point the remaining chain could have been sampled using only the reduced model. Perhaps other approaches to substituting function approximations into expensive likelihood evaluations^76,77^ could provide additional insights to the reduced order modeling approximations.

While we have achieved a significant reduction in computational time with our implementation of the Krylov subspace projection, other model reduction algorithms may yet improve this performance. ^78^ For example, the reduced models considered here achieved levels of accuracy (i.e., relative errors of 10^−5^ or less) that are much higher than one would expect to be necessary to compare models in light of far less accurate data. In light of this finding and the fact that parameter discrimination can be achieved at different levels of accuracy for different combinations of models and data,^79^ we suspect that it could be advantageous to build less accurate models that can be evaluated in less time.

Our present work assumes that the full FSP-based solution can be computed to learn the reduced model bases and to evaluate the second stage likelihood in the ADAMH-FSP-Krylov algorithm. For many problems, the required FSP state space can be so large that it is impossible even to keep the full model in computer memory. Representing the FSP model in a low-rank tensor format^25^ is a promising approach that we plan to investigate in order to overcome this limitation. Another important direction is to extend our work here to models where nonlinear dependence on parameters require the full FSP matrix to be assembled for every parameter evaluation.

Our current work focused on using reduced models for Bayesian estimation of posterior parameters for a given experiment design. In addition to this application, the task of finding optimal parameter fits could also benefit from reduced order modeling. For example, techniques from other engineering fields, such as trust-region methods,^80^ may provide valuable improvements to infer stochastic models from gene expression data. Similar strategies to find reduced FSP models could be utilized to explore sensitivity of single-cell response distributions to parameter variations, which could subsequently be used to compute Fisher Information and design more informative experiments. ^66^ In time, a wealth of algorithms and insights remains to be gained by adapting computational methods from the broader computational science and engineering communities to analyze stochastic gene expression.

## CONCLUSION

There is a clear need for efficient computational algorithms for the uncertainty analysis of gene expression models. In this work, we proposed and investigated new approaches for Bayesian parameter inference of stochastic gene expression parameters from single-cell data. We employed adaptive tuning of proposal distributions in addition to delayed acceptance MCMC and reduced-order modeling. Numerical tests confirmed that the reduced models can be used to significantly speed up the sampling process without incurring much loss in accuracy. While we have only focused on smFISH data in this work, we plan to extend our algorithm to other types of data such as time-course trajectories using time-lapse fluorescence microscopy; for higher-throughput, but lower precision single-cell measurements available using flow cytometry; and eventually for lower throughput but higher content data using single-cell sequencing.

## Supporting Information Available

The proof of the asymptotic convergence of the ADAMH-FSP-Krylov is presented in the supporting information. All MATLAB codes used for this manuscript are open source and available at https://github.com/MunskyGroup/Vo_2019_ADAMHFSP.

## Acknowledgments

Research reported in this publication was supported by the National Institute of General Medical Sciences of the National Institutes of Health under award numbers R25GM105608 and R35GM124747. The work reported here was partially supported by a National Science Foundation grant (DGE-1450032). Any opinions, findings, conclusions or recommendations expressed are those of the authors and do not necessarily reflect the views of the National Science Foundation. The funders had no role in study design, data collection and analysis, decision to publish, or preparation of the manuscript.

## Footnotes

1 https://github.com/lacerbi/multiESS

2 https://github.com/mjlaine/mcmcstat

## References

(1) McAdams, H. H.; Arkin, A. Stochastic mechanisms in gene expression. Proc. Natl. Acad. Sci. U. S. A. 1997, 94, 814–819.

(2) Elowitz, M. B.; Levine, A. J.; Siggia, E. D.; Swain, P. S. Stochastic gene expression in a single cell. Science 2002, 297, 1183–1186.

(3) Kaern, M.; Elston, T. C.; Blake, W. J.; Collins, J. J. Stochasticity in gene expression: from theories to phenotypes. Nature Rev. Genet. 2005, 6, 451–464.

(4) Brehm-Stecher, B. F.; Johnson, E. A. Single-cell microbiology: tools, technologies, and applications. Microbiol. Molecular Biol. Rev. 2004, 68, 538–559.

(5) Schnoerr, D.; Sanguinetti, G.; Grima, R. Approximation and inference methods for stochastic bBiochemical kKinetics - a tutorial review. J. Phys. A 2017, 50.

(6) Femino, A. M.; Fay, F. S.; Fogarty, K.; Singer, R. H. Visualization of single RNA transcripts in situ. Science 1998, 280, 585–590.

(7) Raj, A.; van Oudenaarden, A. Nature, nurture, or chance: stochastic gene expression and its consequences. Cell 2008, 135, 216–226.

(8) Neuert, G.; Munsky, B.; Tan, R. Z.; Teytelman, L.; Khammash, M.; Oudenaarden, A. V. Systematic identification of signal-activated stochastic gene regulation. Science 2013, 339, 584–587.

(9) Gaspar, I.; Ephrussi, A. Strength in numbers: quantitative single-molecule RNA detection assays. 2015, 4, 135–150.

(10) Gomez-Schiavon, M.; Chen, L.; West, A. E.; Buchler, N. E. BayFish: Bayesian Inference of Transcription Dynamics from Population Snapshots of Single-molecule RNA FISH in Single Cells. Genome Biol. 2017, 18, 164.

(11) Gillespie, D. T. A rigorous derivation of the chemical master equation. Physica A 1992, 188, 404–425.

(12) Shepherd, D. P.; Li, N.; Micheva-Viteva, S. N.; Munsky, B.; Hong-Geller, E.; Werner, J. H. Counting small RNA in pathogenic bacteria. Anal. Chem. 2013, 85, 4938–4943.

(13) Munsky, B.; Fox, Z.; Neuert, G. Integrating single-molecule experiments and discrete stochastic models to understand heterogeneous gene transcription dynamics. Methods 2015, 85, 12–21.

(14) Munsky, B.; Khammash, M. The finite state projection algorithm for the solution of the chemical master equation. J. Chem. Phys. 2006, 124, 044104.

(15) Munsky, B.; Li, G.; Fox, Z. R.; Shepherd, D. P.; Neuert, G. Distribution shapes govern the discovery of predictive models for gene regulation. PNAS 2018,

(16) Peherstorfer, B.; Willcox, K.; Gunzburger, M. Survey of multifidelity methods in uncertainty propagation, inference, and optimization. SIAM Review 2018, 60, 550–591.

(17) Asher, M. J.; Croke, B. F. W.; Jakeman, A. J.; Peeters, L. J. M. A review of surrogate models and their application to groundwater modeling. Water Resour. Res. 2015, 51, 5957–5973.

(18) Razavi, S.; Tolson, B. A.; Burn, D. H. Review of surrogate modeling in water resources. Water Resour. Res. 2012, 48.

(19) Schilders, W. H. A.; Vorst, H. A. V. D.; Rommes, J. Eur. Consort. Math. Ind., 1st ed.; Springer-Verlag Berlin Heidelberg, 2008; Vol. 13; pp XI, 471.

(20) Benner, P.; Gugercin, S.; Willcox, K. A survey of model reduction methods for parametric systems. SIAM R 2015, 57, 483–531.

(21) Peleš, S.; Munsky, B.; Khammash, M. Reduction and solution of the chemical master equation using time scale separation and finite state projection. J. Chem. Phys. 2006, 125, 1–13.

(22) Munsky, B.; Khammash, M. The finite state projection approach for the analysis of stochastic noise in gene networks. IEEE Trans. Aut. Contrl. 2008, 53, 201–214.

(23) Tapia, J. J.; Faeder, J. R.; Munsky, B. Adaptive coarse-graining for transient and quasi-equilibrium analyses of stochastic gene regulation. IEEE 51st Conf. Decis. Ctrl. (CDC) 2012, 836, 5361–5366.

(24) Vo, H. D.; Sidje, R. B. Solving the chemical master equation with aggregation and Krylov approximations. 2016, 7093–7098.

(25) Kazeev, V.; Khammash, M.; Nip, M.; Schwab, C. Direct solution of the chemical master equation using quantized tensor trains. PLoS Comput. Biol. 2014, 10.

(26) Dolgov, S.; Khoromskij, B. N. Simultaneous state-time approximation of the chemical master equation using tensor product formats. Numer. Linear Algebra Appl. 2013, 22, 197–219.

(27) Vo, H. D.; Sidje, R. B. An adaptive solution to the chemical master equation using tensors. J. Chem. Phys. 2017, 147.

(28) Dayar, T.; Orhan, M. C. On compact vector formats in the solution of the chemical master equation with backward differentiation. Numer. Linear Algebra Appl. 2018, 25, e2158.

(29) Waldherr, S.; Haasdonk, B. Efficient parametric analysis of the chemical master equation through model order reduction. BMC Sys. Biol. 2012, 6, 81.

(30) Liao, S.; Vejchodský, T.; Erban, R. Tensor methods for parameter estimation and bifurcation analysis of stochastic reaction networks. J. R. Soc. Interface 2015, 12, 20150233.

(31) Oseledets, I. V. Tensor-train decomposition. SIAM J. Sci. Comput. 2011, 33, 2295–2317.

(32) Haario, H.; Saksman, E.; Tamminen, J. An Adaptive Metropolis Algorithm. Bernoulli 2001, 7, 223.

(33) Christen, J. A.; Fox, C. Markov chain Monte Carlo using an approximation. J. Comput. Graph. Stat. 2005, 14, 795–810.

(34) Efendiev, Y.; Hou, T.; Luo, W. Preconditioning Markov chain Monte Carlo simulations using coarse-scale models. SIAM J. Sci. Comput. 2006, 28, 776–803.

(35) Cui, T.; Fox, C.; O’Sullivan, M. J. Bayesian calibration of a large-scale geothermal reservoir model by a new adaptive delayed acceptance Metropolis Hastings algorithm. Water Resources Research 2011, 47.

(36) Cui, T.; Fox, C.; O’Sullivan, M. Adaptive error modelling MCMC sampling for large scale inverse problems. Tech. Report 2011, Fac. of Engr., Univ. of Auckland.

(37) Golightly, A.; Henderson, D. A.; Sherlock, C. Delayed acceptance particle MCMC for exact inference in stochastic kinetic models. Stat. Comput. 2015, 25, 1039–1055.

(38) Saad, Y. Analysis of some Krylov subspace appproximations to the matrix exponential operator. SIAM J. Numer. Anal. 1992, 29, 209–228.

(39) Sidje, R. B. Expokit: A software package for computing matrix exponentials. ACM Trans. Math. Softw. 1998, 24, 130–156.

(40) Burrage, K.; Hegland, M.; MacNamara, S.; Sidje, R. B. In 150th Markov Anniversary Meeting, Charleston, SC, USA; Langville, A., Stewart, W., Eds.; Boson Books, 2006; pp 21–38.

(41) Sidje, R. B.; Vo, H. D. Solving the chemical master equation by a fast adaptive finite state projection based on the stochastic simulation algorithm. Math. Biosci. 2015, 269, 10–16.

(42) Gauckler, L.; Yserentant, H. Regularity and approximability of the solutions to the chemical master equation. ESAIM. Math. Model. 2014, 48, 1757–1775.

(43) Chaturantabut, S.; Sorensen, D. C. Nonlinear model reduction via discrete empirical interpolation. SIAM J. Sci. Comput. 2010, 32, 2737–2764.

(44) Metropolis, N.; Rosenbluth, A. W.; Rosenbluth, M. N.; Teller, A. H.; Teller, E. Equation of state calculations by fast computing machines. J. Chem. Phys. 1953, 21, 1087–1092.

(45) Hastings, W. K. Monte Carlo sampling methods using Markov chains and their applications. Biometrika 1970, 57, 97–109.

(46) Roberts, G. O.; Rosenthal, J. S. General state space Markov chains and MCMC algorithms. Prob. Surv. 2004, 1, 20–71.

(47) Hey, K. L.; Momiji, H.; Featherstone, K.; E Davis, J. R.; H White, M. R.; Rand, D. A.; Finkenst, A. A stochastic transcriptional switch model for single cell imaging data. Biostat. 2015, 16, 655–669.

(48) Tiberi, S.; Walsh, M.; Cavallaro, M.; Hebenstreit, D.; Finkenstädt, B. Bayesian inference on stochastic gene transcription from flow cytometry data. Bioinformatics 2018, 34, i647–i655.

(49) Roberts, G. O.; Gelman, A.; Gilks, W. R. Weak convergence and optimal scaling for random walk Metropolis Hastings algorithms. Annal. Appl. Prob. 1997, 7, 110–120.

(50) Cui, T.; Marzouk, Y. M.; Willcox, K. E. Data-driven model reduction for the Bayesian solution of inverse problems. Int. J. Numer. Meth. Engnr. 2015, 102, 966–990.

(51) Golub, G.; Van Loan, C. Matrix Computations, 4th ed.; John Hopkins University Press, 2012.

(52) Vo, H. D.; Sidje, R. B. Implementation of variable parameters in the Krylov-based finite state projection for solving the chemical master equation. Appl. Math. Comput. 2017, 293, 334–344.

(53) Cao, Y.; Terebus, A.; Liang, J. Accurate chemical master equation solution using multi-finite buffers. Multiscale Model. Simul. 2016, 14, 923–963.

(54) Binev, P.; Cohen, A.; Dahmen, W.; DeVore, R.; Petrova, G.; Wojtaszczyk, P. Convergence rates for greedy algorithms in reduced basis methods. SIAM J. Math. Anal. 2011, 43, 1457–1472.

(55) Roberts, G.; Rosenthal, J. S. Coupling and ergodicity of adaptive Markov chain. J. Appl. Probability 2007, 44, 458–475.

(56) Vats, D.; Flegal, J. M.; Jones, G. L. Multivariate Output Analysis for Markov Chain Monte Carlo. *arXiv* 2017, arXiv:1512.07713v4.

(57) Geweke, J. Evaluating the accuracy of sampling-based approaches to the calculation of posterior moments. Bayesian Statistics. 1992; pp 169–193.

(58) Gillespie, D. T. Exact stochastic simulation of coupled chemical reactions. J. Phys. Chem. 1977, 81, 2340–2361.

(59) Munsky, B.; Neuert, G.; van Oudenaarden, A. Using gene expression noise to understand gene regulation. Science 2012, 336, 183–187.

(60) Peccoud, J.; Ycart, B. Markovian modeling of gene-product synthesis. Theoretical Pop. Biol. 1995, 48, 222 –234.

(61) Golding, I.; Paulsson, J.; Zawilski, S. M.; Cox, E. C. Real-time kinetics of gene activity in individual bacteria. Cell 2005, 123, 1025–1036.

(62) Iyer-Biswas, S.; Hayot, F.; Jayaprakash, C. Stochasticity of gene products from transcriptional pulsing. Phys. Rev. E 2009, 79, 031911.

(63) Gardner, T.; Cantor, C.; Collins, J. Construction of a genetic toggle switch in Escherichia coli. Nature 2000, 403, 339–342.

(64) Tian, T.; Burrage, K. Stochastic models for regulatory networks of the genetic toggle switch. PNAS 2006, 103, 8372–8377.

(65) Munsky, B.; Khammash, M. Identification from stochastic cell-to-cell variation: a genetic switch case study. IET Syst. Biol. 2010, 4, 356–366.

(66) Fox, Z. R.; Munsky, B. The finite state projection based Fisher information matrix approach to estimate information and optimize single-cell experiments. PLOS Comput. Biol. 2019, 15, 1–23.

(67) Golightly, A.; Wilkinson, D. J. Bayesian inference for stochastic kinetic models using a diffusion approximation. Biometrics 2005, 61.

(68) Boys, R. J.; Wilkinson, D. J.; Kirkwood, T. B. Bayesian inference for a discretely observed stochastic kinetic model. Stat. Comput. 2008, 18, 125–135.

(69) Daigle, B. J.; Roh, M. K.; Petzold, L. R.; Niemi, J. Accelerated maximum likelihood parameter estimation for stochastic biochemical systems. BMC Bioinformatics 2012, 13.

(70) Wu, Q.; Smith-Miles, K.; Tian, T. Approximate Bayesian computation schemes for parameter inference of discrete stochastic models using simulated likelihood density. BMC Bioinfo. 2014, 15 Suppl 12, S3.

(71) Andreychenko, A.; Mikeev, L.; Spieler, D.; Wolf, V. Parameter identification for Markov models of biochemical reactions. Computer Aided Verification. Berlin, Heidelberg, 2011; pp 83–98.

(72) Ruess, J.; Lygeros, J. Moment-based methods for parameter inference and experiment design for stochastic biochemical reaction networks. ACM Trans. Model. Comput. Simul. 2015, 25, 8:1–8:25.

(73) Fröhlich, F.; Thomas, P.; Kazeroonian, A.; Theis, F. J.; Grima, R.; Hasenauer, J. Inference for Stochastic Chemical Kinetics Using Moment Equations and System Size Expansion. PLoS Comput. Biol. 2016, 12.

(74) Bonassi, F. V.; You, L.; West, M. Bayesian learning from marginal data in bionetwork models. Stat. Appl. Genet. Mol. Biol. 2011, 10.

(75) Zechner, C.; Ruess, J.; Krenn, P.; Pelet, S.; Peter, M.; Lygeros, J.; Koeppl, H. Moment-based inference predicts bimodality in transient gene expression. PNAS 2012, 109.

(76) Conrad, P. R.; Marzouk, Y. M.; Pillai, N. S.; Smith, A. Accelerating asymptotically exact MCMC for computationally intensive models via local approximations. J. Amer. Stat. Assoc. 2016, 111, 1591–1607.

(77) Conrad, P.; Davis, A.; Marzouk, Y.; Pillai, N.; Smith, A. Parallel local approximation MCMC for expensive models. SIAM/ASA J. Uncertain. 2018, 6, 339–373.

(78) Benner, P.; Cohen, A.; Ohlberger, M.; Wilcox, K. e. Model reduction and approximation: Theory and algorithms; SIAM Publishing, 2017.

(79) Fox, Z.; Neuert, G.; Munsky, B. Finite state projection based bounds to compare chemical master equation models using single-cell data. J. Chem. Phys. 2016, 145.

(80) Qian, E.; Grepl, M.; Veroy, K.; Willcox, K. A certified trust region reduced basis approach to PDE-Constrained optimization. SIAM J. Sci. Comput. 2017,

