## Supplemental Information for "Bayesian estimation for stochastic gene expression using multifidelity models"

### Asymptotic convergence of the ADAMH-FSP-Krylov

#### Preliminaries on adaptive MCMC algorithms

We will derive ergodicity results in the following sections based on Theorem 1 in the paper of Roberts and Rosenthal,[1] and we will use some proof techniques of Theorem 1 from Cui et al.[2] for part of our analysis. All random variables we will discuss below will be of the form $X : \Omega \to \mathcal{X}$ where $\mathcal{X}$ is a metric space with the associated Borel $\sigma$-algebra $B(\mathcal{X})$.

Let $\mathcal{X}$ be the parameter space, assumed to have a metric space topology, and $\pi : B(\mathcal{X}) \to [0,1]$ the target distribution to be sampled from by an adaptive MCMC algorithm. We will assume that $\pi$ has a density $f : \mathcal{X} \to [0, \infty)$. Let $K_\gamma$ denote a transition kernel that depends on an adaptation index $\gamma \in \mathcal{Y}$, and assume that each $K_\gamma$ has $\pi$ as an invariant distribution. We assume that for each fixed $\gamma$, an MCMC algorithm with $K_\gamma$ as the Markov transition kernel will eventually converge to $\pi$, that is

$$\lim_{n\to\infty} \|K^n(x,.) - \pi\|_{TV} = 0$$

where $\|\mu - \nu\|_{TV} = \sup_{B\in B(\mathcal{X})} |\mu(B) - \nu(B)|$ is the total variation distance between two probability measures on $\mathcal{X}$.

Let $X_n$ be the random variable representing the state of the adaptive MCMC at iteration $n$, and let $\Gamma_n$ be the random variable representing the choice of kernel for updating from $X_n$ to $X_{n+1}$. The state of the algorithm is then modeled by the discrete-time stochastic process $\{(X_n, \Gamma_n)\}$, whose transition between steps is determined by the underlying rules of the algorithm. Finally, let $\mathcal{G}_n = \sigma(\{X_0, \ldots, X_n, \Gamma_0, \ldots, \Gamma_n\})$ denote the filtration generated by $\{(X_n, \Gamma_n)\}$. Thus, each $\Gamma_{n+1}$ is a $\mathcal{G}_{n+1}$-measurable random variable.

Roberts and Rosenthal proved the following important result, which gives sufficient conditions for ergodicity of an adaptive MCMC.

**Theorem 1** (Theorem 1 in Roberts and Rosenthal[1]). *Consider an adaptive MCMC algo-*

*rithm with state space $\mathcal{X}$ and adaptation index $\mathcal{Y}$, with transition kernels $K_\gamma,\ \gamma \in \mathcal{Y}$. The algorithm is ergodic if the following conditions hold*

*(i) (Simultaneous uniform ergodicity) For every $\varepsilon > 0$, there exists $N = N(\varepsilon)$ such that*

$$\|K_\gamma^n(x, .) - \pi\|_{TV} < \varepsilon$$

*for every $x \in \mathcal{X}$, $\gamma \in \mathcal{Y}$, and $n > N$.*

*(ii) (Diminishing adaptation) $\lim_{n\to\infty} D_n = 0$ in probability where*

$$D_n = \sup_{x\in\mathcal{X}} \|K_{\Gamma_n}(x, .) - K_{\Gamma_{n+1}}(x, .)\|_{TV}$$

*is a $\mathcal{G}_{n+1}$-measurable random variable.*

We immediately get a useful corollary.

**Corollary 2.** *Consider an adaptive MCMC with state space $\mathcal{X}$ and transition kernels $K_\gamma,\ \gamma \in \mathcal{Y}$ that are ergodic w.r.t $\pi$. Assume that the following conditions are satisfied:*

*(i) The algorithm satisfies the diminishing adaptiation condition.*

*(ii) $\mathcal{X}$ is a compact metric space.*

*(iii) $\mathcal{Y} = \cup_{j=1}^m \mathcal{Y}_j$ where each $\mathcal{Y}_j$ is a compact metric space.*

*(iv) For each $n = 1, 2, \ldots,$ and on each set $\mathcal{X} \times \mathcal{Y}_j$ with the product metric space topology, the mapping*

$$(x, \gamma) \mapsto S(x, \gamma; n, j) = \|K_\gamma^n(x, .) - \pi(.)\|_{TV}$$

*is continuous.*

*Then, the adaptive MCMC algorithm is ergodic.*

*Proof.* Our proof is a modification of the proof of Corollary 3 in.[1] Fix a number $\varepsilon > 0$ and an index $j \in \{1, \ldots, m\}$. Let $W_n^j$ be the set of all $(x, \gamma) \in \mathcal{X} \times \mathcal{Y}_j$ such that

$$S(x, \gamma; n, j) < \varepsilon.$$

Since each kernel is ergodic, for every $(x, \gamma) \in \mathcal{X} \times \mathcal{Y}_j$ there exists some $n$ such that $(x, \gamma) \in W_n^j$, and that $S(x, \gamma; n', j) < \varepsilon$ for all $n' > n$. We thus have

$$\mathcal{X} \times \mathcal{Y}_j = \cup_{n=1}^{\infty} {W_n}^j$$

Due to continuity, each ${W_n}^j$ is an open set. By compactness, there exists a finite subcover $\{{W_{n_i}}^j\}_{i=1}^{r_j}$ for $\mathcal{X} \times \mathcal{Y}_j$. Choose $N_j(\varepsilon)$ to be the maximum of all $n_1, \ldots, n_{r_j}$. Then, choose $N(\varepsilon) = N_1(\varepsilon) + \ldots + N_m(\varepsilon)$, we have $\|K_\gamma^n(x, .) - \pi\|_{TV} < \varepsilon$ for all $n > N(\varepsilon)$ and $(x, \gamma) \in \mathcal{X} \times \mathcal{Y}$. Thus, simultaneous uniform ergodicity is satisfied. Combining with diminishing adaptation, the preceding theorem shows that the algorithm is ergodic. □

#### Convergence of adaptive DAMH with diminishing model adaptations

In this section, we analyze the convergence of an adaptive variant of the DAMH. As seen in the pseudocode of Algorithm 1, this variant modifies the approximation and the proposal density at every step, using the samples accepted so far on the chain. The update of the approximate model occurs randomly, with the upate probability at step $n$ pre-specified as $a(n)$.

**Proposition 3.** *Consider an adaptive delayed acceptance Metropolis-Hastings algorithm with the target distribution supported on a state space $\mathcal{X}$, proposal adaptation space $\mathcal{Y}$, approximation space $\mathcal{Z}$. Let $f$ be the density of the target distribution $\pi$ with respect to a finite reference measure $\lambda$, that is, $\pi(dx) = f(x)\lambda(dx)$. Let $f^\star_{x,\varphi}$ be the family of approximations*

**Algorithm 1** Adaptive Delayed Acceptance MH with probabilistic approximation adaptation.

**Input:**
Target density $f(.)$;
Proposal densities $q_\gamma(.,.)$;
Posterior density approximations $f^\star_{x,\varphi}(.)$;
Adaptation probability $a(n), \quad , n = 1, 2, \ldots$;
Chain length $N$.

Assume that $x_n = x$, $\gamma_n = \gamma$ at iteration $n$.
The next sample is determined by the following steps.

1. Propose a candidate $x'$ from the proposal density $q_\gamma(x, .)$.

2. Compute the first-step acceptance probability

$$\alpha_{\gamma,\varphi}(x, x') = \min\left\{1, \frac{q_\gamma(x', x) f^\star_{x,\varphi}(x')}{q_\gamma(x, x') f^\star_{x,\varphi}(x)}\right\}$$

3. With probability $\alpha_{\gamma,\varphi}$, set $y = x'$. Otherwise, set $y = x$. The actual proposal distribution is

$$Q^\star_{x,\gamma,\varphi}(x, dz) = q_\gamma(x, z)\alpha_{\gamma,\varphi}(x, z)\lambda(dz) + \delta_x(dz)(1 - r_{\gamma,\varphi}(x)),$$

where

$$r_{\gamma,\varphi}(x) = \int_{\mathcal{X}} q_\gamma(x, z')\alpha_{\gamma,\varphi}(x, z')\lambda(dz')$$

is the overall probability that a proposal is accepted in the first step.

4. Set $x_{n+1} = y$ with probability

$$\beta_{\gamma,\varphi}(x, x') = \min\left\{1, \frac{q_\gamma(x', x) f^\star_x(x') f(x')}{q_\gamma(x, x') f^\star_x(x) f(x)}\right\}.$$

Otherwise, set $x_{n+1} = x_n$.

5. With probability $a(n)$, update the approximation $f^\star_{x,\varphi}$.

6. Update the first-step proposal $q_\gamma(x, .)$.

**Output:** Samples $x_1, \ldots, x_N$.

*to $f$. Let $q_\gamma$ be the first-step proposal densities. The algorithm is ergodic under the following conditions:*

*(i)* $\mathcal{X}, \mathcal{Y}$ *are compact metric spaces, and* $\mathcal{Z} = \cup_{j=1}^m \mathcal{Z}_j$ *where each* $\mathcal{Z}_j$ *is a compact metric space.*

*(ii) For each fixed* $\gamma, \varphi$*, the transition kernel* $K_{\gamma,\varphi}$ *is ergodic.*

*(iii)* $\lambda\{x\} = 0$ *for all* $x \in \mathcal{X}$*.*

*(iv) The mapping* $(x, y, \gamma) \mapsto q_\gamma(x, y)$ *is continuous and uniformly bounded on* $\mathcal{X} \times \mathcal{X} \times \mathcal{Y}$ *which is a compact metric space equipped with the product space metric.*

*(v) For each* $y \in \mathcal{Y}$*, the mapping* $(x, \varphi) \mapsto f^\star_{x,\varphi}(y)$ *is continuous on each* $\mathcal{X} \times \mathcal{Z}_j$*.*

*(vi) Diminishing adaptation: The chain* $(\Gamma_n, \Phi_n)$ *satisfies*

$$\lim_{n\to\infty} \sup_{x\in\mathcal{X}} \|K_{\Gamma_{n+1},\Phi_{n+1}}(x, .) - K_{\Gamma_n,\Phi_n}(x, .)\|_{TV} = 0$$

*in probability.*

*Proof.* The ADAMH could be viewed as an adaptive MCMC algorithm with state space $\mathcal{X}$ and adaptation space $\mathcal{Y} \times \mathcal{Z}$. In order to apply corollary 2, we will prove that for any fixed $n = 1, 2, \ldots$, and fixed $j = 1, \ldots, m$, the mapping

$$(x, \gamma, \varphi) \mapsto \|K^n_{\gamma,\varphi}(x, .) - \pi\|_{TV}$$

is continuous on $\mathcal{X} \times \mathcal{Y} \times \mathcal{Z}_j$. In order to do so, we proceed as in the proof of theorem 1 in.[2] Fix $(x, \gamma, \varphi) \in \mathcal{X} \times \mathcal{Y} \times \mathcal{Z}_j$, the transition kernel for the DAMH associated with $(x, \gamma, \varphi)$ is

$$K_{\gamma,\varphi}(x, dz) = q_\gamma(x, z)\alpha_{\gamma,\varphi}(x, z)\beta_{\gamma,\varphi}(x, z)\lambda(dz) + \delta_x(dz)(1 - \rho_{\gamma,\varphi}(x)),$$

where $\alpha_{\gamma,\varphi}(x,z) = \min\left\{1, \frac{q_\gamma(z,x) f^\star_{x,\varphi}(z)}{q_\gamma(x,z) f^\star_{x,\varphi}(x)}\right\}$ is the first step acceptance probability, $\beta_{\gamma,\varphi}(x,z) = \min\left\{1, \frac{q_\gamma(z,x) f^\star_x(z) f(z)}{q_\gamma(x,z) f^\star_x(x) f(x)}\right\}$ is the second step acceptance probability, and

$$\rho_{\gamma,\varphi}(x) = \int_{\mathcal{X}} q_\gamma(x,z)\alpha_{\gamma,\varphi}(x,z)\beta_{\gamma,\varphi}(x,z)\lambda(dz)$$

is the overall probability for a proposal to be accepted.

Fix the value of $z$, then due to conditions (iv) and (v), $g(x,z,\gamma,\varphi) = q_\gamma(x,z)\alpha_{\gamma,\varphi}(x,z)\beta_{\gamma,\varphi}(x,z)$ is jointly continuous in $(x,\gamma,\varphi) \in \mathcal{X} \times \mathcal{Y} \times \mathcal{Z}_j$. Furthermore, condition (iv) implies that the functions $z \mapsto g(x,z,\gamma,\varphi)$ is uniformly bounded for $(x,\gamma,\varphi) \in \mathcal{X} \times \mathcal{Y} \times \mathcal{Z}_j$. By the bounded convergence theorem, $\rho_{\gamma,\varphi}(x)$ is jointly continuous in the three variables $x, \gamma, \varphi$.

By induction, we can show that the $n$-step transition kernel has the form

$$K^n_{\gamma,\varphi}(x,dz) = g_n(x,z,\gamma,\varphi)\lambda(dz) + \delta_x(dz)\left(1 - \rho_{\gamma,\varphi}(x)\right)^n$$

where $g_n$ is an appropriate function that is jointly continuous in $x, \gamma$ and $\varphi$.

From condition (iii), $\delta_x$ and $\pi$ are orthogonal measures. Therefore,

$$\|K^n_{\gamma,\varphi}(x,.) - \pi\|_{TV} = (1 - \rho_{\gamma,\varphi}(x))^n + \frac{1}{2}\int_{\mathcal{X}} (g_n(x,z,\gamma,\varphi) - f(z))\lambda(dz).$$

The integral on the right hand side is jointly continuous in $x, \gamma, \varphi$ due to the bounded convergence theorem. This shows that $\|K^n_{\gamma,\varphi}(x,.) - \pi\|_{TV}$ is continuous in the variable $(x,\gamma,\varphi) \in \mathcal{X} \times \mathcal{Y} \times \mathcal{Z}_j$. From this, conditions (i), (vi) and corollary 2 combined show that the algorithm is ergodic. □

**Proposition 4.** *Assume the ADAMH with probabilistic model adaptation satisfies conditions (i)-(v) in proposition* 3. *Assume further that the proposal is symmetric, that the approximate posterior adaptation probability $a(n) \to 0$ as $n \to \infty$, and that $d_{\mathcal{Y}}(\Gamma_{n+1}, \Gamma_n) \to 0$ in probability (here $d_{\mathcal{Y}}$ denote the metric on $\mathcal{Y}$). Then, the algorithm satisfies diminishing adaptation.*

*Proof.* All conditions for ergodicity in proposition 3 are satisfied, except for the diminishing adaptation that we will verify. Fix a value of $n$. Consider a fixed set of values $(\gamma_j, \varphi_j)_{j=1}^n$ of adaptivity parameters of the ADAMH chain up to iteration $n$.

Fix an event $A \in B(\mathcal{X})$ and $x \in \mathcal{X}$. We have

$$|K_{\gamma_{n+1},\varphi_{n+1}}(x,A) - K_{\gamma_n,\varphi_n}(x,A)| \leq \underbrace{\left|K_{\gamma_{n+1},\varphi_{n+1}}(x,A) - K_{\gamma_n,\varphi_{n+1}}(x,A)\right|}_{D_1} + \underbrace{\left|K_{\gamma_n,\varphi_{n+1}}(x,A) - K_{\gamma_n,\varphi_n}(x,A)\right|}_{D_2}$$

We bound each term separately. First of all, we have $D_2 = 0$ if $\varphi_n = \varphi_{n+1}$ and $D_2 \leq K_{\gamma_n,\varphi_{n+1}}(x,A) + K_{\gamma_n,\varphi_n}(x,A) \leq 2$ if $\varphi_n \neq \varphi_{n+1}$, with the latter event taking place with probability less than $a(n)$.

Due to the symmetry of the proposal, the first and second step acceptance probabilities do not depend on the choice of $\gamma$. This and the uniform continuity of $q_\gamma(x,y)$ gives us

$$\begin{aligned} D_1 &\leq 2\int_{\mathcal{X}} |(q_{\gamma_{n+1}}(x,y) - q_{\gamma_n}(x,y))\alpha_{\varphi_n}(x,y)\beta_{\varphi_n}(x,y)|\lambda(dy) \\ &\leq 2\int_{\mathcal{X}} |q_{\gamma_{n+1}}(x,y) - q_{\gamma_n}(x,y)|\lambda(dy) \\ &\leq C \cdot d_{\mathcal{Y}}(\gamma_n, \gamma_{n+1}) \end{aligned}$$

where $C > 0$ is independent of $x, y, \gamma$ and $\varphi$.

Combining the bounds on $D_1$ and $D_2$ we get

$$|K_{\gamma_{n+1},\varphi_{n+1}}(x,A) - K_{\gamma_n,\varphi_n}(x,A)| \leq C \cdot d_{\mathcal{Y}}(\gamma_n, \gamma_{n+1}) + 2\chi([\varphi_n = \varphi_{n+1}])$$

where $\chi(A) = 1$ if $A$ is true and 0 otherwise. Taking the supremum over all $x$ and $A$ we get

$$D_n = \sup_{x\in\mathcal{X}} \|K_{\gamma_{n+1},\varphi_{n+1}}(x,.) - K_{\gamma_n,\varphi_n}(x,.)\|_{TV} \leq C \cdot d_{\mathcal{Y}}(\gamma_n, \gamma_{n+1}) + 2\chi([\varphi_n \neq \varphi_{n+1}])$$

Fix a scalar $\varepsilon > 0$. The set of runs where $D_n < \varepsilon$ include sample chains where both events $\varphi_n = \varphi_{n+1}$ and $C.d_{\mathcal{Y}}(\gamma_n, \gamma_{n+1}) < \varepsilon$ hold. Therefore, the event $[D_n \geq \varepsilon]$ is a subset of the event $[C \cdot d_{\mathcal{Y}}(\Gamma_n, \Gamma_{n+1}) \geq \varepsilon] \cup [\Phi_n \neq \Phi_{n+1}]$. We therefore have

$$\begin{aligned}\mathbb{P}[D_n \geq \varepsilon] &\leq \mathbb{P}[C \cdot d_{\mathcal{Y}}(\Gamma_n, \Gamma_{n+1}) \geq \varepsilon] + \mathbb{P}[\Phi_n \neq \Phi_{n+1}] \\ &\leq \mathbb{P}[d_{\mathcal{Y}}(\Gamma_n, \Gamma_{n+1}) \geq \varepsilon / C] + a(n).\end{aligned}$$

The last right hand side of the inequality above converges to 0 as $n \to \infty$. Therefore, $D_n$ converges to 0 in probability. The diminishing adaptation condition is satisfied and the algorithm is ergodic. □

#### Regularity of the ROM-based likelihood approximation

Let $\mathcal{S}_j$ be the set of all $n \times j$ matrices $\boldsymbol{Q}$ such that $\boldsymbol{Q}^T\boldsymbol{Q} = \boldsymbol{I}_{j\times j}$. It is known that $\mathcal{S}_j$ with the metric defined by the induced matrix 2-norm

$$\|\boldsymbol{Q}\| = \max_{\boldsymbol{x}\neq \boldsymbol{0}} \frac{\|\boldsymbol{Q}\boldsymbol{x}\|_2}{\|\boldsymbol{x}\|_2}$$

is a compact metric space (indeed, it is the inverse image of $\boldsymbol{I}_{j\times j}$ via the continuous mapping $\boldsymbol{A} \mapsto \boldsymbol{A}^T\boldsymbol{A}$). Let $m_{\max}$ be the maximum dimension allowed in the reduced basis and let $\boldsymbol{\Phi}$ be a particular basis set constructed during a run of the ADAMH chain, then there exists a tuple $(j_1, \ldots, j_{n_B})$ with $1 \leq j_k \leq m_{\max}$ such that

$$\boldsymbol{\Phi} \in \mathcal{S}_{j_1} \times \cdots \times \mathcal{S}_{j_{n_B}} := \boldsymbol{S}_{\boldsymbol{j}}$$

Thus, the set of all possible choices of reduced basis set $\boldsymbol{\Phi}$ is the finite union of all $\boldsymbol{S}_{\boldsymbol{j}}$ with $\boldsymbol{j}$ bounded elementwise by $m_{\max}$. Note that each $\boldsymbol{S}_{\boldsymbol{j}}$ is a compact metric space with the product space topology. Thus, we can apply the theory developed in the previous section to show that the ADAMH-FSP-Krylov is ergodic. The following propositions concern the continuity in the change of the reduced-order approximations with respect to the change in basis.

**Proposition 5.** *Fix a space* $\boldsymbol{S_j}$ *as above, and let* $\boldsymbol{\Phi}$ *and* $\boldsymbol{\Psi}$ *be elements of this space. For for every fixed* $\boldsymbol{\theta} \in \Theta$ *we have*

$$L^{\star}_{\boldsymbol{\Psi}}(\boldsymbol{\theta}) \to L^{\star}_{\boldsymbol{\Phi}}(\boldsymbol{\theta})$$

*as* $\boldsymbol{\Psi} \to \boldsymbol{\Phi}$ *in* $\boldsymbol{S_j}$*, where* $L^{\star}_{\boldsymbol{\Phi}}$ *is the approximation to the FSP log-likelihood as defined in eq.* (16) *of the main text.*

*Proof.* From eq. (9) of the main text, it is clear that the mapping $\boldsymbol{\Phi} \mapsto p_{\boldsymbol{\Phi}}(t_k)$ is continuous on $\boldsymbol{S_j}$ for all time points $t_k$. The mapping $\boldsymbol{\Phi} \mapsto L^{\star}_{\boldsymbol{\Phi}}(\boldsymbol{\theta})$ is a composition of continuous mappings $\boldsymbol{\Phi} \mapsto p_{\boldsymbol{\Phi}}(t_k)$ and $\boldsymbol{p} \mapsto \sum_{j=1}^{n_i} \log(\varepsilon \vee \boldsymbol{p}_j)$ and is therefore continuous. □

#### Ergodicity of the ADAMH-FSP-Krylov algorithm

**Proposition 6.** *The ADAMH-FSP-Krylov algorithm is ergodic.*

*Proof.* We apply proposition 3 with $\mathcal{X} = \boldsymbol{\Theta}$. The proposal densities of the first step are Gaussian with $\gamma$ being the modified empirical covariance matrix as in the adaptive Metropolis Algorithm.[3] Similar to the proof of Theorem 1 in Haario et al.,[3] we can take $\mathcal{Y}$ to be a closed, bounded subset of the set of positive definite matrices. The reduced model space is $\mathcal{Z} = \cup_{\boldsymbol{j}} \boldsymbol{S_j}$ the finite union of the compact spaces $\boldsymbol{S_j}$ with $\boldsymbol{j} \leq m_{\max}$ pointwise. These spaces satisfy condition (i), and the proposal density satisfies condition (iv).

The posterior density is

$$f(\boldsymbol{\theta}) = \pi_0(\boldsymbol{\theta}) \exp(-L(D|\boldsymbol{\theta})),$$

and the approximate posterior densities are

$$f^{\star}_{\boldsymbol{\Phi}}(\boldsymbol{\theta}) = \pi_0(\boldsymbol{\theta}) \exp(-L^{\star}_{\boldsymbol{\Phi}}(D|\boldsymbol{\theta})),$$

where these are the densities of the true and approximate posterior distributions with respect to the Lebesgue measure. From Theorem 1 in Christen and Fox,[4] condition (ii) is satisfied.

Condition (v) is then satisfied using proposition 5.

Since the empirical covariances are computed from values in a bounded set, the modification to the empirical covariance matrix $\gamma$ at step $n$ is $O(1/n)$, so changes in $\Gamma_n$ converge to 0 (see Haario et al.[3]). Thus, the conditions in proposition 4 are satisfied. The algorithm therefore satisfies all sufficient conditions for ergodicity outlined in proposition 3. □

### References


(1) Roberts, G.; Rosenthal, J. S. Coupling and ergodicity of adaptive Markov chain. *J. Appl. Probability* **2007**, *44*, 458–475.

(2) Cui, T.; Fox, C.; O'Sullivan, M. Adaptive error modelling MCMC sampling for large scale inverse problems. *Tech. Report* **2011**, *Fac. of Engr., Univ. of Auckland*.

(3) Haario, H.; Saksman, E.; Tamminen, J. An Adaptive Metropolis Algorithm. *Bernoulli* **2001**, *7*, 223.

(4) Christen, J. A.; Fox, C. Markov chain Monte Carlo using an approximation. *J. Comput. Graph. Stat.* **2005**, *14*, 795–810.
